# Deletion or inhibition of PTPRO mitigates diet-induced hepatic steatosis and inflammation in obesity

**DOI:** 10.1101/2022.09.05.506586

**Authors:** Takafumi Shintani, Ryoko Suzuki, Yasushi Takeuchi, Takuji Shirasawa, Masaharu Noda

## Abstract

Chronic inflammation plays crucial roles in obesity-induced metabolic diseases. We herein demonstrated that mice lacking the protein tyrosine phosphatase receptor type O (PTPRO) exhibited the hyper-obese phenotype when fed a high-fat/high-sucrose diet. However, *Ptpro*-KO mice with hyperobesity showed the markedly small accumulation of ectopic fat in the liver, improvements in lipid and glucose homeostasis, and low-grade systemic inflammation associated with low macrophage activation. Expression of *protein tyrosine phosphatase 1b* (*Ptp1b*), an enzyme which is known to be implicated in metabolic disorders, was also suppressed in *Ptpro*-KO mice. The administration of AKB9778, a specific inhibitor of PTPRO, to highly obese *ob*/*ob* mice reproduced the phenotypes of *Ptpro*-KO mice along with the amelioration of inflammation. We revealed that an increase in the phosphorylation of Tyr(117) in vimentin, a component of intermediate filaments, by the inhibition of PTPRO promoted the growth of lipid droplets in adipocytes. The improvement in metabolic conditions with the attenuation of inflammation in *Ptpro*-KO mice was explained by the low activation of NFκb, a key transcription factor for inflammatory response, in adipose tissue. This is the first study to show that PTPRO is a promising target to ameliorate hepatic steatosis and metabolic disorders.

## INTRODUCTION

Two billion adults worldwide are classified as overweight, 650 million of whom are obese (World Health Organization, 2016). Obesity is a well-established risk factor for a myriad of conditions affecting multiple body systems, including cardiovascular diseases, diabetes, and even cancers (Hotamisligil, 2017). One of the mechanisms by which obesity causes metabolic complications is the accumulation of fatty acids in non-adipose tissues, such as the liver, skeletal muscle, and pancreas. The accumulation of ectopic fat in non-adipose tissues is induced by fatty acids that leak from adipose tissue when storage exceeds the capacity limitation, resulting in dysfunctional adipose and non-adipose tissues (Goossens, 2017; Longo et al., 2019). Accumulating evidence has suggested that metabolic alterations allowing adipose tissue to grow beyond the upper limit of the normal range are associated with the negligible accumulation of ectopic lipids in other organs, such as the liver (Abreu-Vieira et al., 2015; Kim et al., 2008; Tanaka et al., 2014; Verschoor et al., 2021). For example, transgenic *ob*/*ob* mice lacking leptin while overexpressing adiponectin in adipose tissue showed higher insulin sensitivity and better glucose metabolism in addition to a markedly larger adipose tissue mass than control *ob*/*ob* mice (Kim et al., 2008).

Adipose tissue is well recognized as a significant depot of stromal cells, including immune cells, which exert detrimental effects on overall health (Kahn et al., 2019; Suganami et al., 2012). Obesity-induced inflammation stems from expanded adipose tissue and then spreads to other tissues, including the liver and muscle, resulting in low-grade, but extensive systemic inflammation (Kahn et al., 2019; Suganami et al., 2012). The mechanism by which obesity induces chronic inflammation appears to be a key factor for characterizing the pathogenic cascades of obesity-related diseases. Recent studies demonstrated that exposure to free fatty acids promoted proinflammatory M1-polarized macrophage phenotypes through the activation of the Toll-like receptor (TLR) 4 complex (Lee et al., 2001; Shi et al., 2006; Suganami et al., 2007). M1 macrophages secrete pro-inflammatory cytokines, such as tumor necrosis factor α (TNFα), interleukin-6 (IL6), and monocyte chemoattractant protein-1 (MCP-1), and promote
 the infiltration of other macrophages into adipose tissue (Kahn et al., 2019; Suganami et al., 2012). Adipocytes also secrete bioactive substances called adipokines, including leptin, adiponectin, TNFα, and IL6, which regulate energy homeostasis and inflammation in the body (Goossens et al., 2017; Kahn et al., 2019; Suganami et al., 2012). Numerous studies have shown that the obesity-induced inflammation of adipose tissue leads to the dysregulation of adipokine production. Therefore, in inflamed adipose tissue, macrophages and adipocytes both actively produce pro-inflammatory factors that directly impair systemic insulin sensitivity, which, in turn, induces the development of hepatic steatosis, hypertriglyceridemia, and type 2 diabetes mellitus (T2DM) (Goossens et al., 2017; Kahn et al., 2019; Suganami et al., 2012).

We previously demonstrated at the cellular level that members of the same R3 subfamily of receptor-like protein tyrosine phosphatases (RPTPs), consisting of PTPRO, PTPRJ, PTPRB, and PTPRH, individually regulate the activities of specific receptor-type protein tyrosine kinases (PTKs), including the insulin receptor, through dephosphorylation (Sakuraba et al., 2013; Shintani et al., 2006; Shintani et al., 2015; Shintani et al., 2017). PTPRJ-deficient mice showed improved insulin sensitivity and leptin sensitivity *in vivo* (Shintani et al., 2015; Shintani et al., 2017). However, the role of PTPRO in metabolic regulation remains largely unknown. In the present study, we found that *Ptpro*-KO mice fed a high-fat/high-sucrose diet (HFHSD) exhibited a markedly larger body mass than wild-type (WT) mice, but had a healthy liver. In these *Ptpro*-KO mice, the accumulation of ectopic fat in the liver was markedly small, and inflammation in the liver and adipose tissue was significantly suppressed. Moreover, the administration of a specific inhibitor of PTPRO to *ob*/*ob* mice induced adipose tissue expansion together with a decrease in the accumulation of ectopic fat in the liver, reduced inflammation, and improved insulin sensitivity. Therefore, PTPRO appears to be involved in the mechanisms regulating the limitation of fat accumulation in adipose tissue and inducing inflammation in obesity.

## RESULTS

### *Ptpro*-KO mice are protected from hepatic steatosis induced by a high-fat diet

Western blotting and immunostaining analyses showed that *Ptpro*-KO mice are indeed deficient in the expression of PTPRO proteins (Figure 1-figure supplement 1). We measured body weight gain and food intake in the growing period of WT and *Ptpro*-KO mice, and found no significant differences between the two genotypes when fed a normal diet (ND) (Figure 1-figure supplement 2A). However, under HFHSD feeding, *Ptpro*-KO mice showed continuous weight gain, in contrast to WT mice which showed a slowdown in weight gain after 20 weeks of age (Figure 1A, left). During the period when weight differences were observed between the two genotypes, food intake by *Ptpro*-KO mice was significantly higher than that by WT mice (Figure 1A, right). At the age of 40 weeks, the average weight of WT mice was ~50 g, while that of *Ptpro*-KO mice increased to >70 g, a severely obese state (Figure 1B). Since male and female *Ptpro*-KO mice showed a similar overweight phenotype under HFHSD (Compare Figure 1A and Figure 1-figure supplement 2B), male mice were used in the present study.

**Figure 1.**
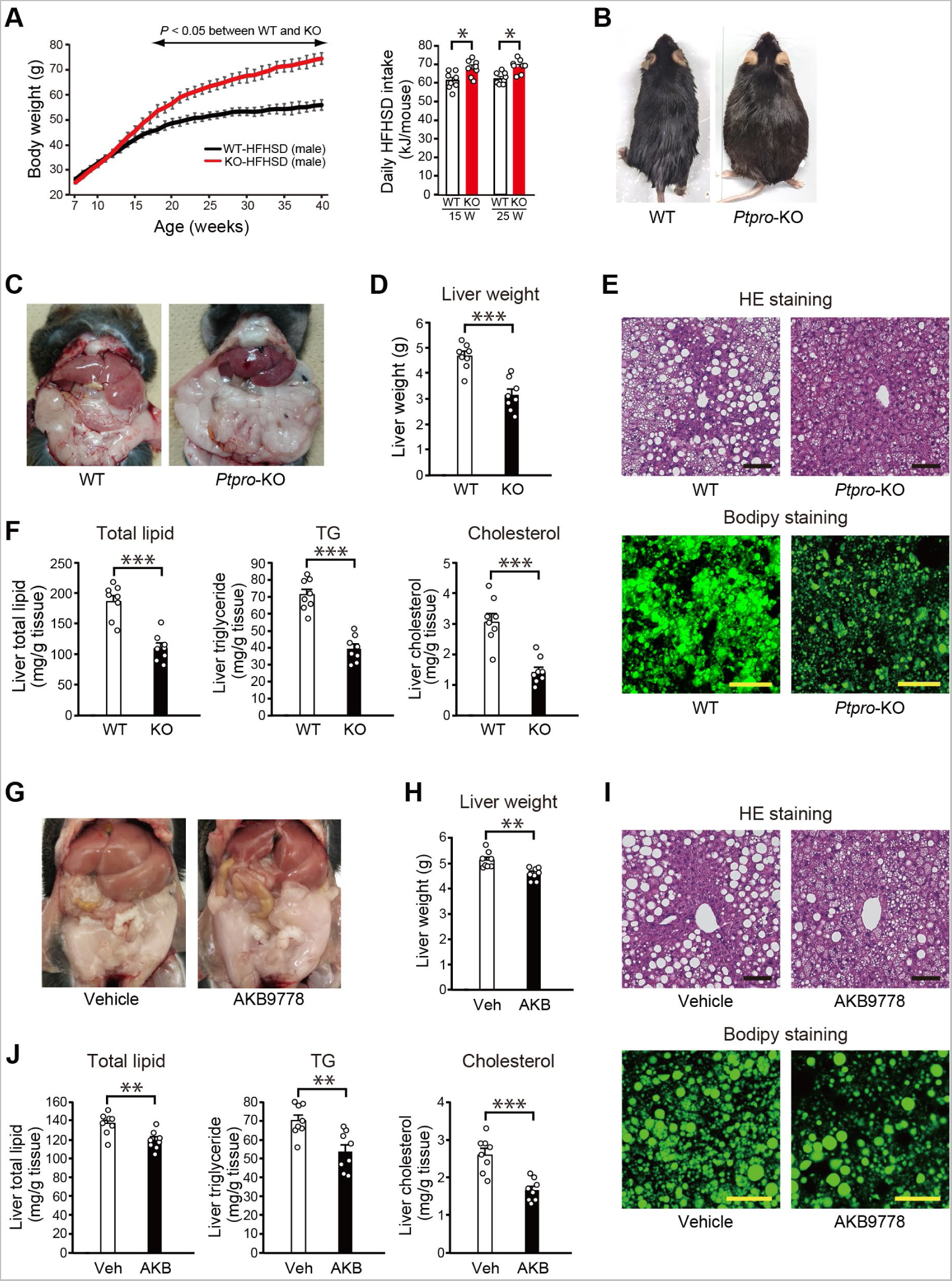
Increased body weight gain without hepatic steatosis in *Ptpro*-KO mice and the amelioration of hepatic steatosis by the AKB9778 treatment in *ob*/*ob* mice. (A) Weekly body weight gains in wild-type (WT) and *Ptpro*-KO (KO) male mice fed HFHSD (left). Daily HFHSD intake by WT and *Ptpro*-KO male mice at 15 and 25 weeks of age (right). n = 8 each. (B) Representative appearance of HFHSD-fed WT and *Ptpro*-KO mice at 40 weeks of age. (C) Representative abdominal views of HFHSD-fed WT and *Ptpro*-KO mice at 24 weeks of age. (D) Liver weight of HFHSD-fed WT and *Ptpro*-KO mice at 24 weeks of age. (E) HE and Bodipy staining of liver sections of HFHSD-fed WT and *Ptpro*-KO mice at 24 weeks of age. (F) Total lipid, triglyceride (TG), and cholesterol levels in the livers of HFHSD-fed WT and *Ptpro*-KO mice at 24 weeks of age. (G) Representative abdominal views of ND-fed *ob*/*ob* mice treated with vehicle (Veh) or AKB9778 (AKB) for 4 weeks from 16 weeks of age. (H) Liver weight of *ob*/*ob* mice treated with vehicle or AKB9778. (I) HE and Bodipy stainings of liver sections of *ob*/*ob* mice treated with vehicle or AKB9778. Bars: 100 μm. (J) Total lipid, triglyceride (TG), and cholesterol levels in the livers of *ob*/*ob* mice treated with vehicle or AKB9778. Values are means ± SEM (n = 8 each). *P* values are based on the unpaired Student’s *t*-test. NS, not significant; \**P* < 0.05, \*\**P* < 0.01, \*\*\**P* < 0.001. **Source data 1.** Weekly body weight in WT and *Ptpro*-KO male mice fed HFHSD, and daily HFHSD intake by WT and *Ptpro*-KO male mice at 15 and 25 weeks of age (Figure 1A). **Source data 2.** Liver weight of WT and *Ptpro*-KO mice fed HFHSD (Figure 1D). **Source data 3.** Total lipid, triglyceride (TG), and cholesterol levels in the livers of HFHSD-fed WT and *Ptpro*-KO mice (Figure 1F). **Source data 4.** Liver weight of *ob*/*ob* mice treated with vehicle or AKB9778 (Figure 1H). **Source data 5.** Total lipid, triglyceride (TG), and cholesterol levels in the livers of *ob*/*ob* mice treated with vehicle or AKB9778 (Figure 1J). **Figure supplement 1.** Absence of PTPRO proteins in *Ptpro*-KO mice. **Figure supplement 2.** Body weight gain and food intake in *Ptpro*-KO mice fed ND or HFHSD. **Figure supplement 3.** Expression of PTPRO in white adipose tissue. **Figure supplement 4.** Dose-response curves of the inhibitory activity of AKB9778 against different PTPs.

In contrast to the livers of WT mice with obesity that turned yellowish due to fatty liver, those of *Ptpro*-KO mice with hyper obesity showed a normal red appearance under HFHSD at 24 weeks of age (Figure 1C). The liver weight of *Ptpro*-KO mice was ~70% that of WT mice (WT; 4.68 ± 0.18 g: KO; 3.14 ± 0.22 g: *P* < 0.0001) (Figure 1D). HE staining of paraffin sections and Bodipy FL staining of frozen sections revealed that lipid accumulation in the liver was markedly suppressed in *Ptpro*-KO mice (Figure 1E); accordingly, biochemical analyses demonstrated that the amounts of lipids (total lipids, triglycerides (TG), and cholesterol) that accumulated in the liver were significantly lower in *Ptpro*-KO mice than in WT mice (Figure 1F). These results indicate that hepatic steatosis induced by HFHSD feeding was markedly attenuated in *Ptpro*-KO mice despite their hyper obesity.

Expression analyses of *Ptpro* mRNA using real-time PCR revealed that *Ptpro* mRNA was highly expressed in white adipose tissue (WAT) among several tissues, particularly visceral WAT (such as epididymal adipose tissue) (Figure 1-figure supplement 3A). On the other hand, *Ptpro* mRNA expressed in subcutaneous WAT was relatively low. When the differentiation process of 3T3-L1 preadipocytes was examined *in vitro*, the expression level of *Ptpro* was found to increase with adipocyte maturation along with the growth of lipid droplets (Figure 1-figure supplement 3B). Furthermore, immunohistochemical analyses indicated that PTPRO proteins were expressed not only in differentiated adipocytes, but also in proinflammatory M1 macrophages in adipose tissue (Figure 1-figure supplement 3C). These results suggest multiple roles for PTPRO in the lipid accumulation in adipocytes as well as in the inflammation of adipose tissue.

### The administration of AKB9778, an inhibitor of PTPRO, improves the liver condition in genetically obese (*ob/ob)* mice, similar to PTPRO-deficient mice

A chemical compound termed AKB9778 was originally developed as an inhibitor of PTPRB in 2013, which belongs to the R3 subfamily of RPTPs as PTPRO (Goel et al., 2013). AKB9778 has been shown to inhibit angiogenesis in the retina and choroid due to the suppression of PTPRB during development (Shen et al., 2014). To estimate the inhibitory activity of AKB9778 on other members of the R3 subfamily of RPTPs, we assessed IC_50_ values by an *in vitro* analysis using the artificial fluorescent substrate, DiFMUP. AKB9778 exhibited high inhibitory activities for PTPRO and PTPRJ as well as PTPRB (Figure 1-figure supplement 4); the highest activity was against PTPRO (IC_50_ = 0.21 nM) among the members tested (IC_50_ values for PTPRJ and PTPRB were 1.14 and 2.58 nM, respectively).

We intraperitoneally administered AKB9778 (10 mg/kg body weight) daily to leptin-deficient *ob*/*ob* mice, a model of severe obesity under ND feeding, for 4 weeks from 16 weeks of age. No significant differences were observed in body weight gains between the AKB9778-treated group and vehicle group during this period. The final body weight of the AKB9778-treated group (58.90 ± 0.88 g) was similar to that of the vehicle group (58.64 ± 0.51 g) at 20 weeks of age. The visible appearance of the liver in the vehicle group was obviously white, whereas that of the AKB9778-treated group apparently improved at 20 weeks of age (Figure 1G). Liver weight was significantly lower in the AKB9778-treated group than in the vehicle group (AKB9778; 4.66 ± 0.08 g: Vehicle; 5.14 ± 0.11 g: *P* = 0.0017) (Figure 1H). HE and Bodipy FL stainings of liver sections both indicated that the accumulation of lipids in the liver was significantly reduced by the AKB9778 treatment (Figure 1I). Quantitative biochemical analyses also showed that total lipid, TG, and cholesterol levels in the liver were all significantly lower in the AKB9778-treated group than in the control group (Figure 1J). These changes induced by the AKB9778 treatment were consistent with those observed in *Ptpro*-KO mice.

We then compared the mRNA expression levels of inflammatory markers in the livers of *Ptpro*-KO mice and AKB9778-treated *ob*/*ob* mice with those in the respective control mice by quantitative PCR. No significant differences were detected in the mRNA expression levels of *Tnfa*, *IL6*, or *Mcp1* between WT and *Ptpro*-KO mice with a lean appearance under ND (Figure 2A). In contrast, the expression levels of *Tnfa*, *IL6*, and *Mcp1* all markedly increased in WT mice fed HFHSD; however, these increases were strongly suppressed in *Ptpro*-KO mice (Figure 2A). The mRNA expression levels of *Tnfa*, *IL6*, and *Mcp1* were consistently lower in the AKB9778-treated group than in the vehicle group in *ob*/*ob* mice (Figure 2B). These results indicated that the inflammatory response in the liver induced by the ectopic accumulation of lipids was strongly suppressed in obese *Ptpro*-KO mice and AKB9778-treated *ob*/*ob* mice.

**Figure 2.**
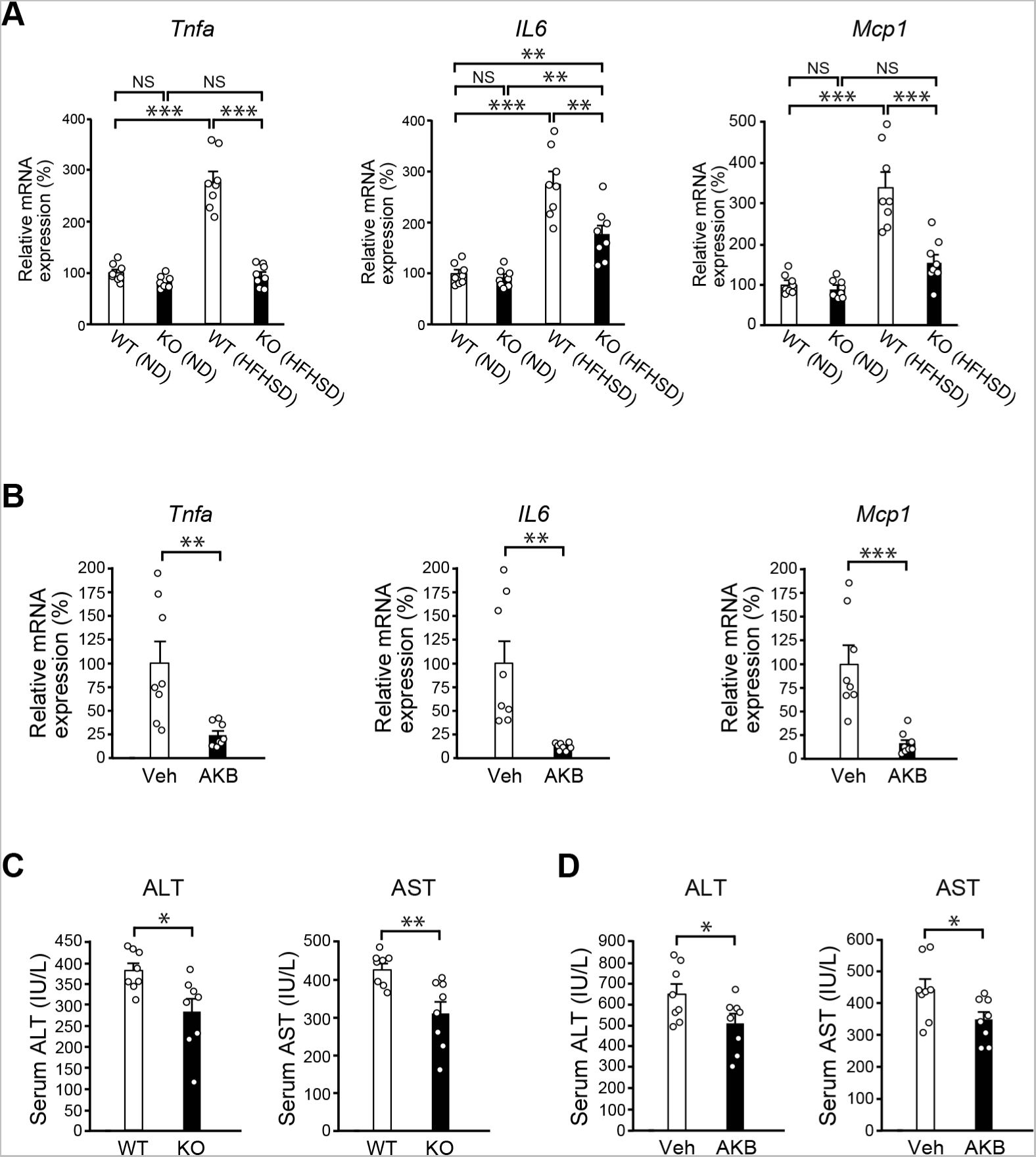
Reduced hepatic inflammation in *Ptpro*-KO mice and AKB9778-treated *ob*/*ob* mice. (A) Expression levels of mRNAs for *Tnfa*, *IL6*, and *Mcp1* in the livers of WT and *Ptpro*-KO (KO) mice under ND- or HFHSD-fed conditions at 24 weeks of age. Data are normalized to the expression level of *Gapdh*, and shown as relative values. Values are means ± SEM (n = 8 each). *P* values are based on a one-way ANOVA followed by Tukey’s test. (B) Expression levels of mRNAs for *Tnfa*, *IL6*, and *Mcp1* in the livers of ND-fed *ob*/*ob* mice treated with vehicle (Veh) or AKB9778 (AKB) for 4 weeks from 16 weeks of age. (C) Serum levels of ALT and AST in WT and *Ptpro*-KO (KO) mice fed HFHSD at 24 weeks of age. Values are means ± SEM (n = 8 each). *P* values are based on the unpaired Student’s *t*-test. (D) Serum levels of ALT and AST in *ob*/*ob* mice after the treatment with vehicle (Veh) or AKB9778 (AKB) for 4 weeks. Values are means ± SEM (n = 8 each). *P* values are based on the unpaired Student’s *t*-test. NS, not significant; \**P* < 0.05, \*\**P* < 0.01, \*\*\**P* < 0.001. **Source data 1.** Expression levels of mRNAs for *Tnfa, IL6*, and *Mcp1* in the livers of WT and *Ptpro*-KO mice under ND- or HFHSD-fed conditions (Figure 2A). **Source data 2.** Expression levels of mRNAs for *Tnfa, IL6*, and *Mcp1* in the livers of ND-fed *ob*/*ob* mice treated with vehicle or AKB9778 (Figure 2B). **Source data 3.** Serum levels of ALT and AST in WT and *Ptpro*-KO mice fed HFHSD (Figure 2C). **Source data 4.** Serum levels of ALT and AST in *ob*/*ob* mice after the treatment with vehicle or AKB9778 (Figure 2D).

The activities of alanine aminotransferase (ALT) and aspartate aminotransferase (AST), markers of liver damage, were significantly lower in HFHSD-fed *Ptpro*-KO mice than in HFHSD-fed WT mice (*P* = 0.0152 and *P* = 0.0045, respectively) (Figure 2C). Consistently, the AKB9778-treated group of *ob*/*ob* mice also showed significantly lower ALT and AST activities than the vehicle group (*P* = 0.0457 and *P* = 0.0308, respectively) (Figure 2D). Therefore, the liver phenotypes induced by obesity were significantly suppressed by the *Ptpro* deficiency and AKB9778 treatment, suggesting a crucial role for PTPRO activity in the induction of hepatic steatosis and inflammation in obesity.

### The *Ptpro* deficiency and AKB9778 treatment induce the expansion of adipose tissue

In contrast to the liver, the weights of adipose tissues, including epididymal adipose tissue, were ~45% higher on average in *Ptpro*-KO mice than in WT mice at 24 weeks of age under HFHSD (*P* = 0.0006) (Figure 3A and Table 1); the weights of individual adipose tissues in *Ptpro*-KO mice (epididymal, 2.96 ± 0.19 g; mesenteric 1.32 ± 0.03 g; retroperitoneal, 2.52 ± 0.06 g; subcutaneous, 7.76 ± 0.21 g; n = 8) were markedly higher than those in WT mice (epididymal, 2.06 ± 0.09 g; mesenteric, 0.96 ± 0.03 g; retroperitoneal, 1.74 ± 0.07 g; subcutaneous, 5.09 ± 0.21 g; n = 8). The ratio of the fat mass to the body weight of each part was significantly higher in *Ptpro*-KO mice than in WT mice (Table 1). In paraffin sections of epididymal adipose tissue, *Ptpro*-KO mice had a higher number of larger adipocytes than WT mice; the average diameter of adipocytes in *Ptpro*-KO mice was ~1.3-fold larger than those in WT mice (*P* < 0.0001) (Figure 3B). Consistently, the mass of all adipose tissues increased by ~20% on average in *ob*/*ob* mice treated with AKB9778 (Figure 3C and Table 2), along with the appearance of larger adipocytes than in the vehicle *ob*/*ob* group (Figure 3D). These results suggested that the ectopic lipid in the liver was transferred to adipose tissue by the treatment with AKB9778.

**Figure 3.**
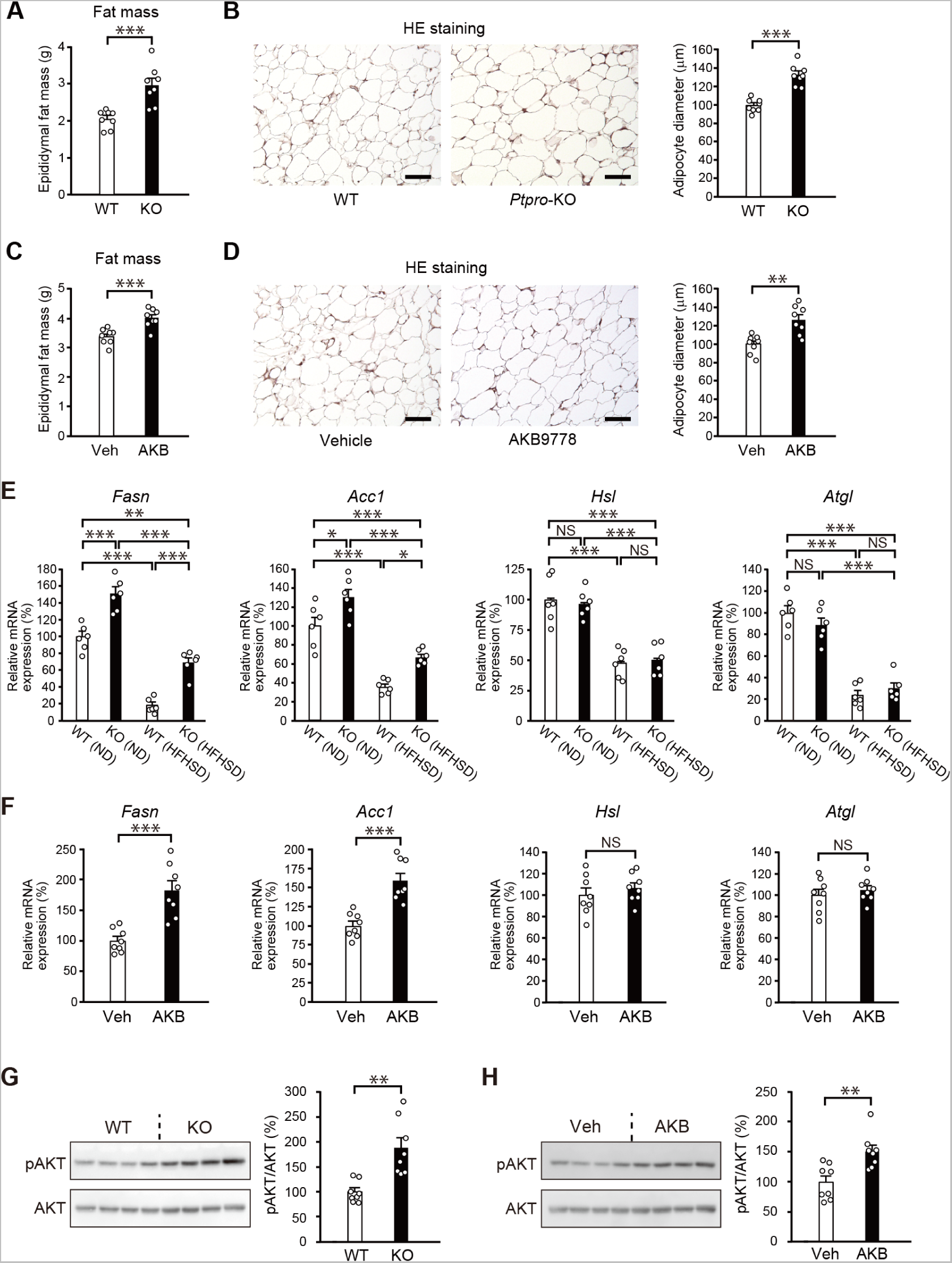
Expansion of adipose tissue in *Ptpro*-KO mice and AKB9778-treated *ob*/*ob* mice. (A) Epididymal fat weight of HFHSD-fed WT and *Ptpro*-KO mice at 24 weeks of age. n = 8 each. (B) HE staining of epididymal fat sections of WT and *Ptpro*-KO mice fed as above. Bars: 100 μm. The right graph shows quantitative analyses of adipocyte diameters (n = 8 each). (C) Epididymal fat weight of ND-fed *ob*/*ob* mice treated with vehicle or AKB9778 for 4 weeks. n = 8 each. (D) HE staining of epididymal fat sections of *ob*/*ob* mice treated with vehicle or AKB9778. Bars: 100 μm. The right graph shows quantitative analyses of adipocyte diameters (n = 8 each). (E) Expression levels of mRNAs for *Fasn*, *Acc1*, *Hsl*, and *Atgl* in the epididymal adipose tissue of WT and *Ptpro*-KO (KO) mice fed ND or HFHSD. Data are normalized to the expression level of *Gapdh*, and shown as relative values. n = 6 each. (F) Expression levels of mRNAs for *Fasn*, *Acc1*, *Hsl*, and *Atgl* in the epididymal adipose tissue of *ob*/*ob* mice treated with vehicle (Veh) or AKB9778 (AKB) for 4 weeks. Data are normalized to the expression level of *Gapdh*, and shown as relative values. n = 8 each. (G) Western blot analyses of pAKT and total AKT in epididymal fat lysates prepared from HFHSD-fed WT and *Ptpro*-KO mice at 24 weeks of age. The right graph shows quantitative analyses, which represents the relative ratio of pAKT/AKT (n = 8 each). (H) Western blot analyses of pAKT and total AKT in epididymal fat lysates prepared from *ob*/*ob* mice treated with vehicle or AKB9778 for 4 weeks. The right graph shows quantitative analyses, which represent the relative ratio of pAKT/AKT (n = 8 each). Values are means ± SEM. *P* values are based on the unpaired Student’s *t*-test (A-D, F-H) or a one-way ANOVA followed by Tukey’s test (E). NS, not significant; \**P* < 0.05, \*\**P* < 0.01, \*\*\**P* < 0.001. **Source data 1.** Epididymal fat weight of HFHSD-fed WT and *Ptpro*-KO mice (Figure 3A). **Source data 2.** Adipocyte diameter of HFHSD-fed WT and *Ptpro*-KO mice (Figure 3B). **Source data 3.** Epididymal fat weight of *ob*/*ob* mice treated with vehicle or AKB9778 (Figure 3C). **Source data 4.** Adipocyte diameter of *ob*/*ob* mice treated with vehicle or AKB9778 (Figure 3D). **Source data 5.** Expression levels of mRNAs for *Fasn, Acc1, Hsl,* and *Atgl* in the epididymal adipose tissue of WT and *Ptpro*-KO (Figure 3E). **Source data 6.** Expression levels of mRNAs for *Fasn, Acc1, Hsl,* and *Atgl* in the epididymal adipose tissue of *ob*/*ob* mice treated with vehicle or AKB9778 (Figure 3F). **Source data 7.** Western blot analyses of pAKT and total AKT in epididymal fat lysates prepared from HFHSD-fed WT and *Ptpro*-KO mice (Figure 3G). **Source data 8.** Western blot analyses of pAKT and total AKT in epididymal fat lysates prepared from *ob*/*ob* mice treated with vehicle or AKB9778 (Figure 3H).

**Table 1:** Body weight and fat weight of HFHSD-fed WT and *Ptpro*-KO mice at 24 weeks of age.

|  | WT | KO |  |
| --- | --- | --- | --- |
| Body weight | 50.88 ± 0.76 | 62.77 ± 1.39 | (*P* < 0.0001) |
| Epididymal fat (g) | 2.05 ± 0.09 | 2.96 ± 0.19 | (*P* = 0.0006) |
| Ratio to BW (%) | 4.03 ± 0.14 | 4.69 ± 0.20 | (*P* = 0.0176) |
| Mesenteric fat (g) | 0.96 ± 0.03 | 1.32 ± 0.03 | (*P* < 0.0001) |
| Ratio to BW (%) | 1.89 ± 0.05 | 2.10 ± 0.03 | (*P* = 0.0032) |
| Retroperitoneal fat (g) | 1.74 ± 0.08 | 2.52 ± 0.06 | (*P* < 0.0001) |
| Ratio to BW (%) | 3.41 ± 0.12 | 4.01 ± 0.08 | (*P* = 0.0009) |
| Subcutaneous fat (g) | 5.09 ± 0.21 | 7.76 ± 0.21 | (*P* < 0.0001) |
| Ratio to BW (%) | 10.03 ± 0.43 | 12.41 ± 0.45 | (*P* = 0.0019) |

Values are means ± SEM. *P* values are based on unpaired Student's t-test.
**Source data 1.** Body weight and fat weight of HFHSD-fed WT and *Ptpro*-KO mice.

**Table 2:**
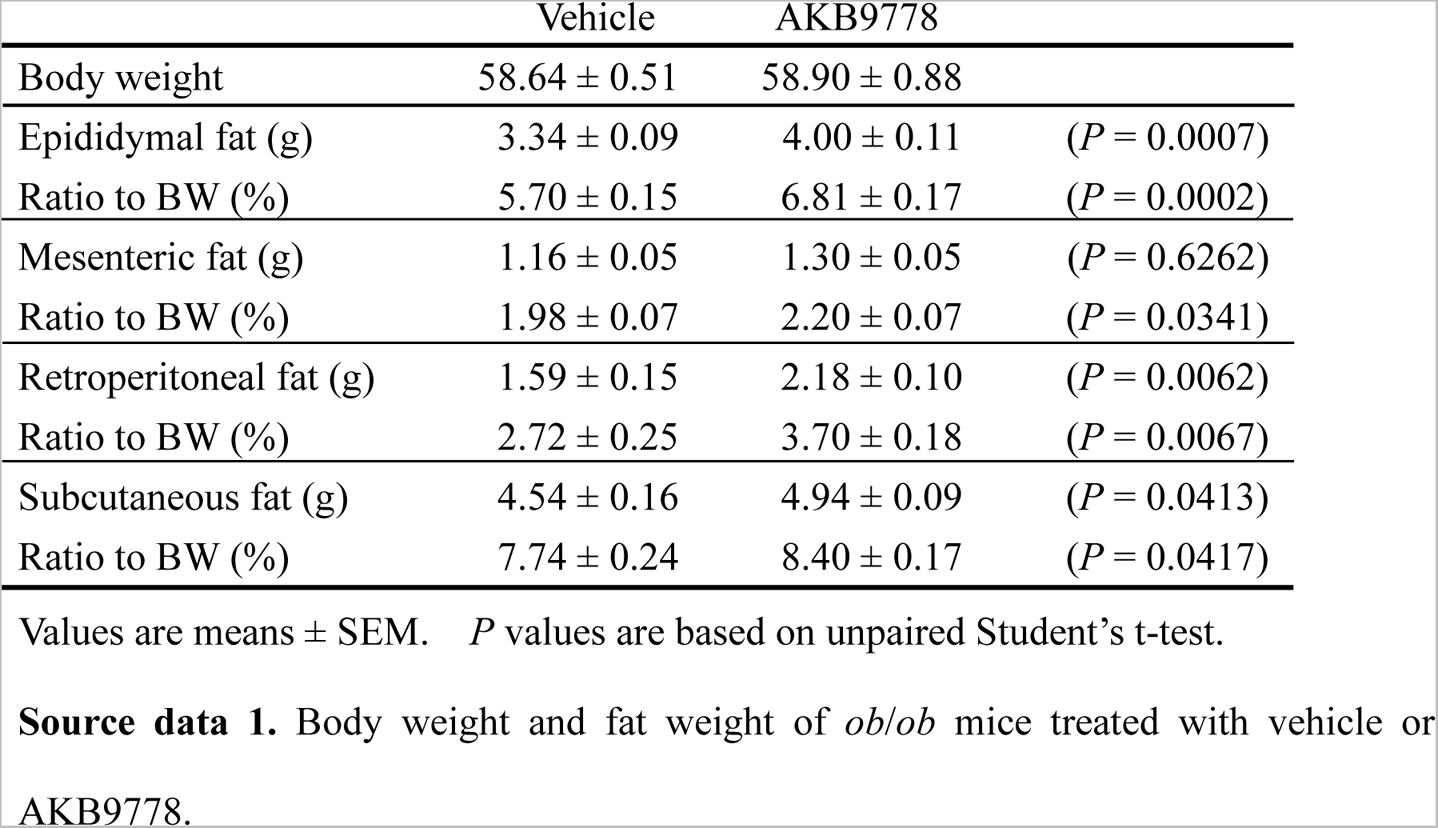
Body weight and fat weight of *ob*/*ob* mice treated with vehicle or AKB9778 for 4 weeks.

|  | Vehicle | AKB9778 |  |
| --- | --- | --- | --- |
| Body weight | 58.64 ± 0.51 | 58.90 ± 0.88 |  |
| Epididymal fat (g) | 3.34 ± 0.09 | 4.00 ± 0.11 | (*P* = 0.0007) |
| Ratio to BW (%) | 5.70 ± 0.15 | 6.81 ± 0.17 | (*P* = 0.0002) |
| Mesenteric fat (g) | 1.16 ± 0.05 | 1.30 ± 0.05 | (*P* = 0.6262) |
| Ratio to BW (%) | 1.98 ± 0.07 | 2.20 ± 0.07 | (*P* = 0.0341) |
| Retroperitoneal fat (g) | 1.59 ± 0.15 | 2.18 ± 0.10 | (*P* = 0.0062) |
| Ratio to BW (%) | 2.72 ± 0.25 | 3.70 ± 0.18 | (*P* = 0.0067) |
| Subcutaneous fat (g) | 4.54 ± 0.16 | 4.94 ± 0.09 | (*P* = 0.0413) |
| Ratio to BW (%) | 7.74 ± 0.24 | 8.40 ± 0.17 | (*P* = 0.0417) |

Values are means ± SEM. *P* values are based on unpaired Student's t-test.
**Source data 1.** Body weight and fat weight of *ob/ob* mice treated with vehicle or AKB9778.

To characterize alterations in gene expression in adipose tissue in *Ptpro*-KO mice and AKB9778-treated mice, we examined genes related to lipid metabolism in adipose tissue. The expression levels of the lipogenic genes *fatty acid synthase* (*Fasn*) and *acetyl-CoA carboxylase 1* (*Acc1*) were higher in *Ptpro*-KO mice than in WT mice under ND feeding conditions (*P* < 0.0001 and *P* = 0.0192, respectively) (Figure 3E). Under HFHSD feeding conditions, the expression levels of these genes in adipose tissue were significantly suppressed in both genotypes (Figure 3E); however, they markedly remained higher in *Ptpro*-KO mice than in WT mice (*P* < 0.0001 and *P* = 0.0159, respectively). On the other hand, no significant differences were observed in the expression levels of the lipase genes, *hormone-sensitive lipase* (*Hsl*) and *adipose triglyceride lipase* (*Atgl*), in adipose tissues between WT and *Ptpro*-KO mice irrespective of whether they were fed ND or HFHSD (Figure 3E). The expression levels of *Fasn* and *Acc1* were consistently higher in AKB9778-treated *ob*/*ob* mice than in vehicle mice (*P* = 0.0002 and *P* = 0.0001, respectively), while the expression levels of *Hsl* and *Atgl* did not significantly differ between the two groups (Figure 3F). These results indicate that the expression of lipogenic genes, but not lipolytic genes, in adipose tissue were specifically and negatively controlled by PTPRO.

We also examined the activation of AKT (protein kinase B) in HFHSD-fed mice because PI3-kinase/AKT signaling is known to promote lipogenesis in adipose tissue (Huang et al., 2018). The activation of AKT (estimated by the phosphorylation levels of Ser-473 of AKT) was stronger in *Ptpro*-KO mice than in WT mice (*P* = 0.0052) (Figure 3G). The pAKT/AKT ratio was consistently higher in AKB9778-treated *ob*/*ob* mice than in vehicle mice (*P* = 0.0034) (Figure 3H). These results again indicated that the *Ptpro* deficiency and AKB9778 treatment promoted lipogenesis in adipose tissue by activating PI3K/AKT signaling.

### The *Ptpro* deficiency and AKB9778 treatment suppress inflammation in adipose tissue

The expansion of adipose tissue is generally associated with tissue inflammation (Goossens, 2017; Lee et al., 2001; Shi et al., 2006). Therefore, we examined the levels of inflammatory marker genes in epididymal adipose tissue by quantitative PCR. Under ND conditions, the gene expression of the inflammatory marker, *Tnfa* was significantly lower in *Ptpro*-KO mice than in WT mice (*P* = 0.0088) (Figure 4A). HFHSD feeding significantly increased the expression levels of the inflammatory markers *Tnfa*, *IL6*, and *Mcp1* in WT mice, but not in *Ptpro*-KO mice (*P* < 0.0001 each) (Figure 4A). The suppression of *Tnfa*, *IL6*, and *Mcp11* was also observed in the adipose tissue of AKB9778-treated *ob*/*ob* mice (*P* = 0.0059, *P* = 0.0174, and *P* = 0.0281, respectively) (Figure 4B). The expression of protein tyrosine phosphatase 1B (PTP1B) in adipose tissue has been reported to be increased by inflammation induced by obesity (Nieto-Vazquez et al., 2008; Zabolotny et al., 2008). Consistent with these reports, HFHSD feeding significantly increased the expression levels of *Ptp1b* in WT mice, but not in *Ptpro*-KO mice (*P* = 0.0002) (Figure 4A). The suppression of *Ptp1b* was also observed in the adipose tissue of AKB9778-treated *ob*/*ob* mice (P < 0.0001) (Figure 4B).

**Figure 4.**
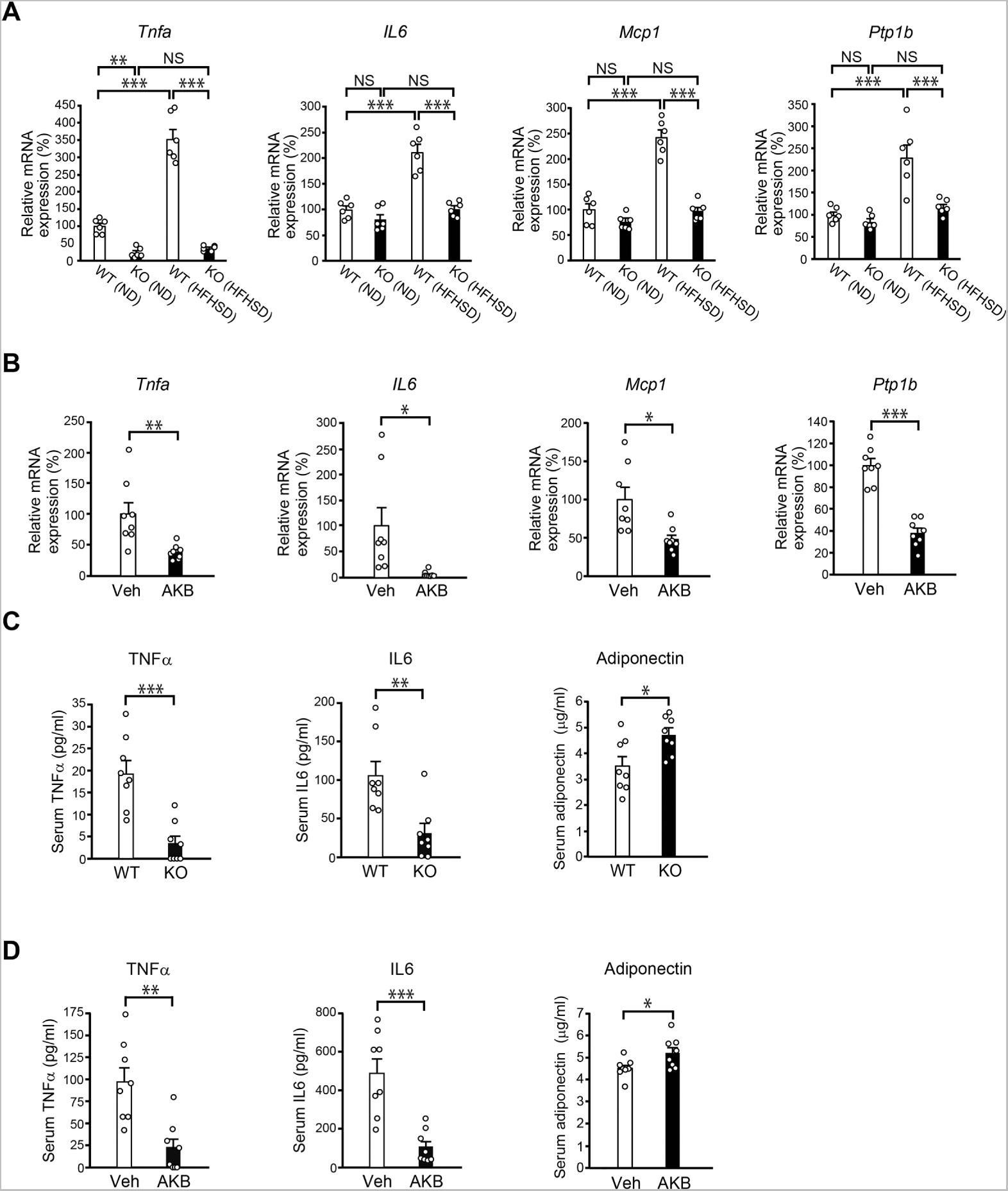
Reduced expression of pro-inflammation factors in *Ptpro*-KO mice and AKB9778-treated *ob*/*ob* mice. (A) Expression levels of mRNAs for *Tnfa*, *IL6*, *Mcp-1*, and *Ptp1b* in the epididymal adipose tissue of WT and *Ptpro*-KO (KO) mice fed ND or HFHSD at 24 weeks of age. Data are normalized to the expression level of *Gapdh*, and shown as relative values. n = 6 each. (B) Expression levels of mRNAs for *Tnfa*, *IL6*, *Mcp-1*, and *Ptp1b* in the epididymal adipose tissue of ND-fed *ob*/*ob* mice after the treatment with vehicle (Veh) or AKB9778 (AKB) for 4 weeks from 16 weeks of age. Data are normalized to the expression level of *Gapdh* and shown as relative values. n = 8 each. (C) Serum levels of TNFα, IL6, and adiponectin in HFHSD-fed WT and *Ptpro*-KO mice. n = 8 each. (D) Serum levels of TNFα, IL6, and adiponectin in *ob*/*ob* mice treated with vehicle or AKB9778 for 4 weeks. n = 8 each. Values are means ± SEM. *P* values are based on a one-way ANOVA followed by Tukey’s test (A) or the unpaired Student’s *t*-test (B-D). NS, not significant; \**P* < 0.05, \*\**P* < 0.01, \*\*\**P* < 0.001. **Source data 1.** Expression levels of mRNAs for *Tnfa, IL6, Mcp-1*, and *Ptp1b* in the epididymal adipose tissue of WT and *Ptpro*-KO mice (Figure 4A). **Source data 2.** Expression levels of mRNAs for *Tnfa, IL6, Mcp-1*, and *Ptp1b* in the epididymal adipose tissue of *ob*/*ob* mice treated with vehicle or AKB9778 (Figure 4B). **Source data 3.** Serum levels of TNFα, IL6, and adiponectin in HFHSD-fed WT and *Ptpro*-KO mice (Figure 4C). **Source data 4.** Serum levels of TNFα, IL6, and adiponectin in *ob*/*ob* mice treated with vehicle or AKB9778 (Figure 4D). **Figure supplement 1.** Serum leptin levels in HFHSD-fed WT and *Ptpro*-KO mice.

Furthermore, the serum levels of TNFα and IL6 were markedly lower in obese *Ptpro*-KO mice (*P* = 0.0003 and *P* = 0.0029, respectively) and AKB9778-treated *ob*/*ob* mice (*P* = 0.0015 and *P* = 0.0003, respectively) than in the respective control groups (Figure 4C, D). On the other hand, serum levels of adiponectin, an adipokine that exerts anti-inflammatory effects, were markedly higher in *Ptpro*-KO mice than in WT mice (*P* = 0.0175) (Figure 4C), whereas serum levels of leptin, which exhibits proinflammatory activity, did not significantly differ between the two genotypes (Figure 4-figure supplement 1). Similarly, serum adiponectin levels were higher in the AKB9778-treated group than in the vehicle group (*P* = 0.0357) (Figure 4D). Therefore, obese *Ptpro*-KO mice and AKB9778-treated *ob*/*ob* mice showed significantly low inflammation in adipose tissues and the systemic level.

The strong induction of *CD11c*, the M1 macrophage marker, and *F4/80*, the mature macrophage marker, by HFHSD feeding in WT mice was similarly suppressed in *Ptpro*-KO mice (*P* < 0.0001 each) (Figure 5A) and AKB9778-treated mice (*P* = 0.0459 and *P* = 0.0034, respectively) (Figure 5B). The immunostaining of epididymal adipose tissue with an anti-F4/80 antibody represents the accumulation of mature macrophages, which is visible as a brown-colored area. In the lean state with ND, F4/80 signals in adipose tissue were almost negligible in WT and *Ptpro*-KO mice (Figure 5-figure supplement 1). Consistent with the results of qPCR, the F4/80 signal was significantly lower in *Ptpro*-KO mice than in WT mice under HFHSD feeding conditions (*P* = 0.006) (Figure 5C), indicating that the infiltration of macrophages into adipose tissue was low in *Ptpro*-KO mice: The number of crown-like structures with F4/80 signals, which are formed by inflammatory M1 macrophages around dying adipocytes, was consistently low in *Ptpro*-KO mice (Figure 5C). Furthermore, the F4/80-positive area in epididymal adipose tissue was smaller in AKB9778-treated *ob*/*ob* mice (*P* = 0.0106) (Figure 5D).

**Figure 5.**
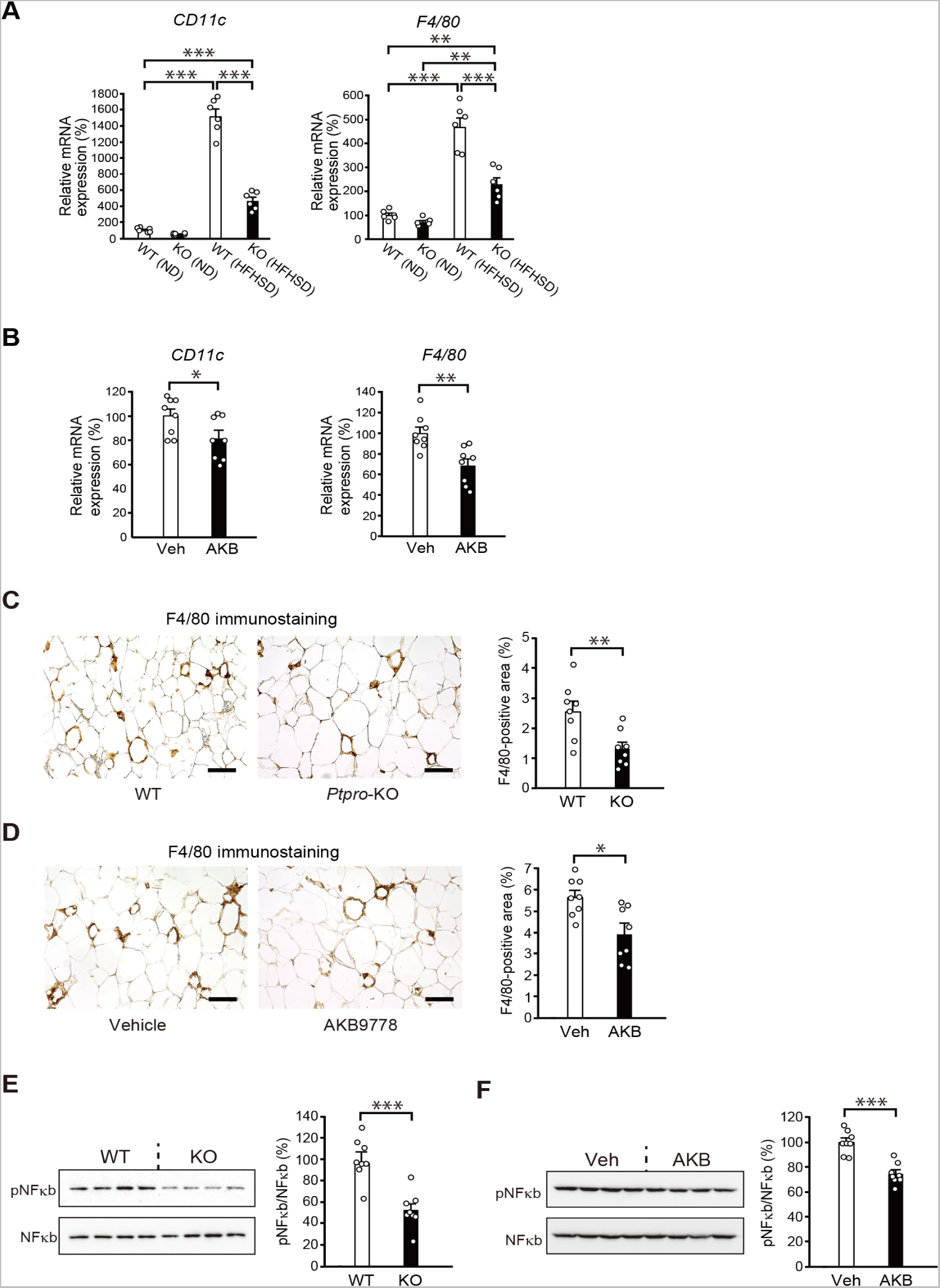
Reduced accumulation and activation of macrophages in the adipose tissue of *Ptpro*-KO mice and AKB9778-treated *ob*/*ob* mice. (A) Expression levels of mRNAs for *Cd11c* and *F4/80* in the epididymal adipose tissue of WT and *Ptpro*-KO (KO) mice fed ND or HFHSD at 24 weeks of age. Data are normalized to the expression level of *Gapdh*, and shown as relative values. n = 6 each. (B) Expression levels of mRNAs for *Cd11c* and *F4/80* in the epididymal adipose tissue of ND-fed *ob*/*ob* mice after the treatment with vehicle (Veh) or AKB9778 (AKB) for 4 weeks from 16 weeks of age. Data are normalized to the expression level of *Gapdh* and shown as relative values. n = 8 each. (C) F4/80 immunostaining of the epididymal fat sections of WT and *Ptpro*-KO mice fed HFHSD. Bars: 100 μm. The right graph shows quantitative analyses of F4/80-positive areas (n = 8 each). (D) F4/80 immunostaining of epididymal fat sections of *ob*/*ob* mice treated with vehicle or AKB9778 for 4 weeks. Bars: 100 μm. The right graph shows a summary of quantitative analyses of F4/80-positive areas (n = 8 each). (E) Western blot analyses of pNFκb and total NFκb in the epididymal fat lysates of HFHSD-fed WT and *Ptpro*-KO mice. The right graphs show quantitative analyses, which represent the relative ratio of pNFκb/NFκb. (F) Western blot analyses of pNFκb and total NFκb in the epididymal fat lysates of *ob*/*ob* mice treated with vehicle or AKB9778 for 4 weeks. The right graphs show quantitative analyses, which represent the relative ratio of pNFκb/NFκb. Values are means ± SEM. *P* values are based on a one-way ANOVA followed by Tukey’s test (A) or the unpaired Student’s *t*-test (B-F). \**P* < 0.05, \*\**P* < 0.01, \*\*\**P* < 0.001. **Source data 1.** Expression levels of mRNAs for *Cd11c* and *F4/80* in the epididymal adipose tissue of WT and *Ptpro*-KO mice (Figure 5A). **Source data 2.** Expression levels of mRNAs for *Cd11c* and *F4/80* in the epididymal adipose tissue of *ob*/*ob* mice treated with vehicle or AKB9778 (Figure 5B). **Source data 3.** F4/80 immunostaining of the epididymal fat sections of WT and *Ptpro*-KO mice fed HFHSD (Figure 5C). **Source data 4.** F4/80 immunostaining of the epididymal fat sections of *ob*/*ob* mice treated with vehicle or AKB9778 (Figure 5D). **Source data 5.** Western blot analyses of pNFκb and total NFκb in the epididymal fat lysates of HFHSD-fed WT and *Ptpro*-KO mice (Figure 5E). **Source data 6.** Western blot analyses of pNFκb and total NFκb in the epididymal fat lysates of *ob*/*ob* mice treated with vehicle or AKB9778 (Figure 5F). **Figure supplement 1.** F4/80 immunostaining of the adipose tissue sections of ND-fed WT and *Ptpro*-KO mice.

We also examined the activation of the transcriptional factor NFκb, which is responsible for the initiation of inflammatory responses in a large number of tissues (Kahn et al., 2019; Napetschnig, 2013). The pNFκb (active form of NFκb)/NFκb ratio was markedly lower in *Ptpro*-KO mice than in WT mice in epididymal adipose tissue (*P* = 0.0002) (Figure 5E), whereas the expression levels of NFκb proteins did not significantly differ between the two genotypes (Figure 5E). This result indicated that the activation of NFκb was suppressed in *Ptpro*-KO mice. The suppressed activation of NFκb was also observed in adipose tissue of AKB9778-treated *ob*/*ob* mice (*P* < 0.0001) (Figure 5F). The attenuation of inflammation in adipose tissue in obese *Ptpro*-KO and AKB9778-treated *ob*/*ob* mice suggested that the activity of PTPRO was involved in the activation of NFκb, leading to the induction of inflammation. The result showing that the expression of PTPRO increases with adipocyte maturation associated with the growth of lipid droplets (Figure 1-figure supplement 3B) is consistent with the fact that inflammation in adipose tissue occurs at the end stage of lipid accumulation when PTPRO activity is high.

### The *Ptpro* deficiency and AKB9778 treatment suppress adipose tissue fibrosis

Persistent inflammatory stress in adipose tissue often induces fibrosis, which is characterized by the excessive deposition of extracellular matrix (ECM) components (Sun, et al., 2013; Marcelin et al., 2019). Tissue fibrosis also produces severe impairment in adipose tissue function. Transforming growth factor β (TGFβ) plays a pivotal role in adipose tissue fibrosis by increasing the expression of matrix components, such as collagen and fibronectin, and inhibitors of matrix metalloproteases (Marcelin et al., 2019). We found that HFHSD feeding markedly increased the expression of *Tgfb1* in the adipose tissue of WT mice, but not *Ptpro*-KO mice; the expression of *Tgfb1* in *Ptpro*-KO mice fed HFHSD remained at the same level as that in WT mice fed ND (Figure 6A). The mRNA expression levels of *collagens* (*Col1a* and *Col3a*) and *tissue inhibitor of matrix metalloproteases 1* (*TIMP-1*) were also increased by HFHSD feeding in WT mice; however, these increases were markedly attenuated in *Ptpro*-KO mice (Figure 6A). The expression levels of *Col1a*, *Col3a*, and *TIMP-1* were consistently lower in AKB9778-treated *ob*/*ob* mice (Figure 6B) than in vehicle mice, except for *Tgfb1* (Figure 6B).

**Figure 6.**
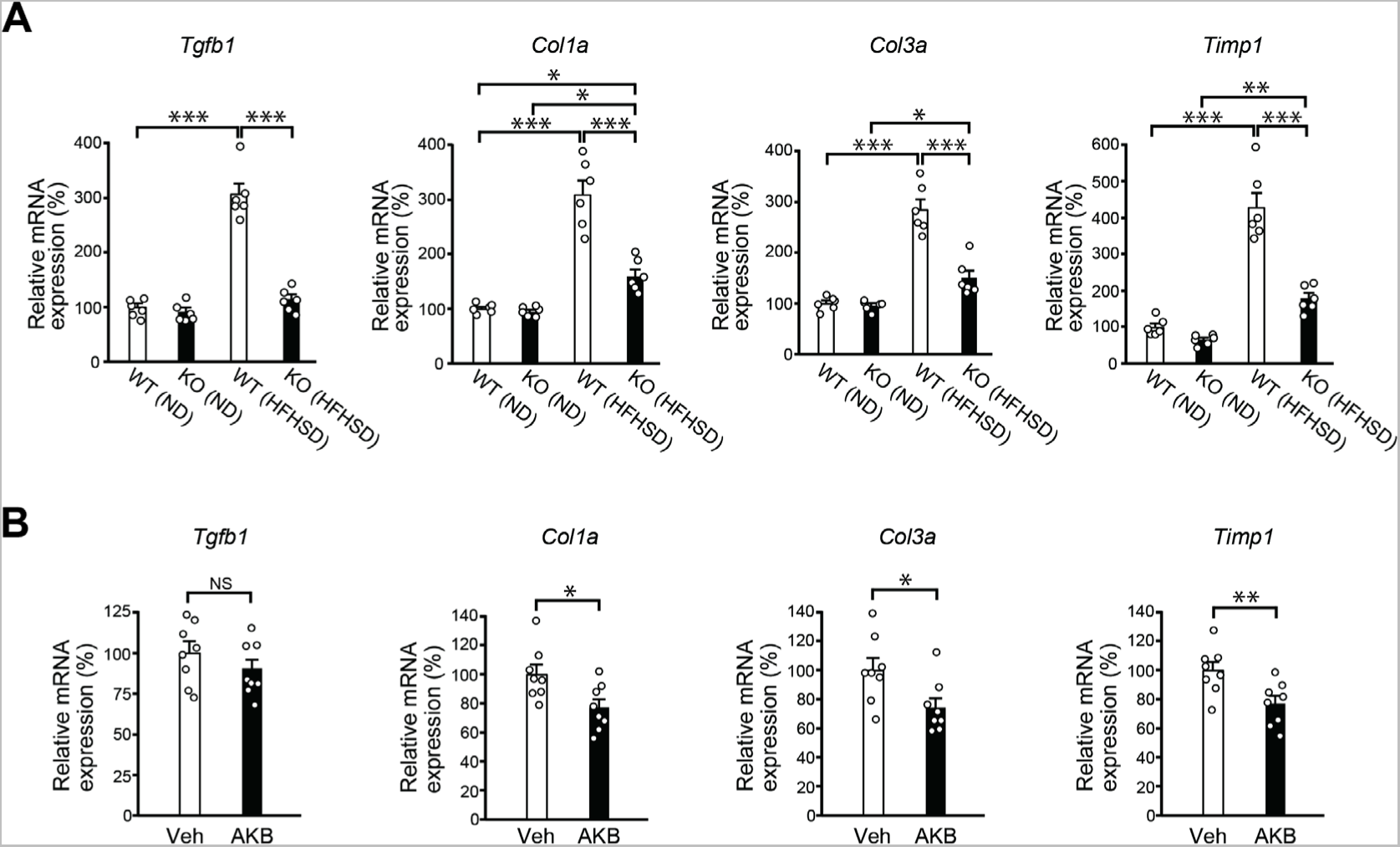
Reduced expression of fibrosis-related genes in the epididymal adipose tissue of *Ptpro*-KO mice and AKB9778-treated *ob*/*ob* mice. (A) mRNA expression levels of *Tgfb1*, *Col1a*, *Col3a*, and *Timp1* in the epididymal adipose tissue of WT and *Ptpro*-KO (KO) mice fed HFHSD at 24 weeks of age. Data are normalized to the expression level of *Gapdh* and shown as relative values. Values are means ± SEM (n = 6 each). *P* values are based on a one-way ANOVA followed by Tukey’s test. (B) mRNA expression levels of *Tgfb1, Col1a*, *Col3a*, and *Timp1* in the epididymal adipose tissue of ND-fed *ob*/*ob* mice treated with vehicle (Veh) or AKB9778 (AKB) for 4 weeks. Data are normalized to the expression level of *Gapdh* and shown as relative values. Values are means ± SEM (n = 8 each). *P* values are based on the unpaired Student’s *t*-test. NS, not significant; \**P* < 0.05, \*\**P* < 0.01, \*\*\**P* < 0.001. **Source data 1.** Expression levels of mRNAs for *Tgfb1, Col1a, Col3a*, and *Timp1* in the epididymal adipose tissue of WT and *Ptpro*-KO mice (Figure 6A). **Source data 2.** Expression levels of mRNAs for *Tgfb1, Col1a, Col3a*, and *Timp1* in the epididymal adipose tissue of *ob*/*ob* mice treated with vehicle or AKB9778.

Masson’s trichrome and Sirius Red staining techniques, which detect collagen fibers, also revealed markedly less fibrosis in *Ptpro*-KO mice than in WT mice (Figure 7A and B, upper). In addition, the positive area of α-smooth muscle actin (α-SMA), a biomarker of activated fibroblasts, was markedly smaller in *Ptpro*-KO mice than in WT mice (Figure 7C, upper). Not only Masson’s trichrome staining (Figure 7A, lower) and Sirius Red staining (Figure 7B, lower), but also α-SMA immunohistostaining (Figure 7C, lower) showed that their positive areas were smaller in AKB9778-treated *ob*/*ob* mice than in the control group. These results indicated that the *Ptpro* deficiency and AKB9778 treatment suppressed adipose tissue fibrosis induced by HFHSD feeding.

**Figure 7.**
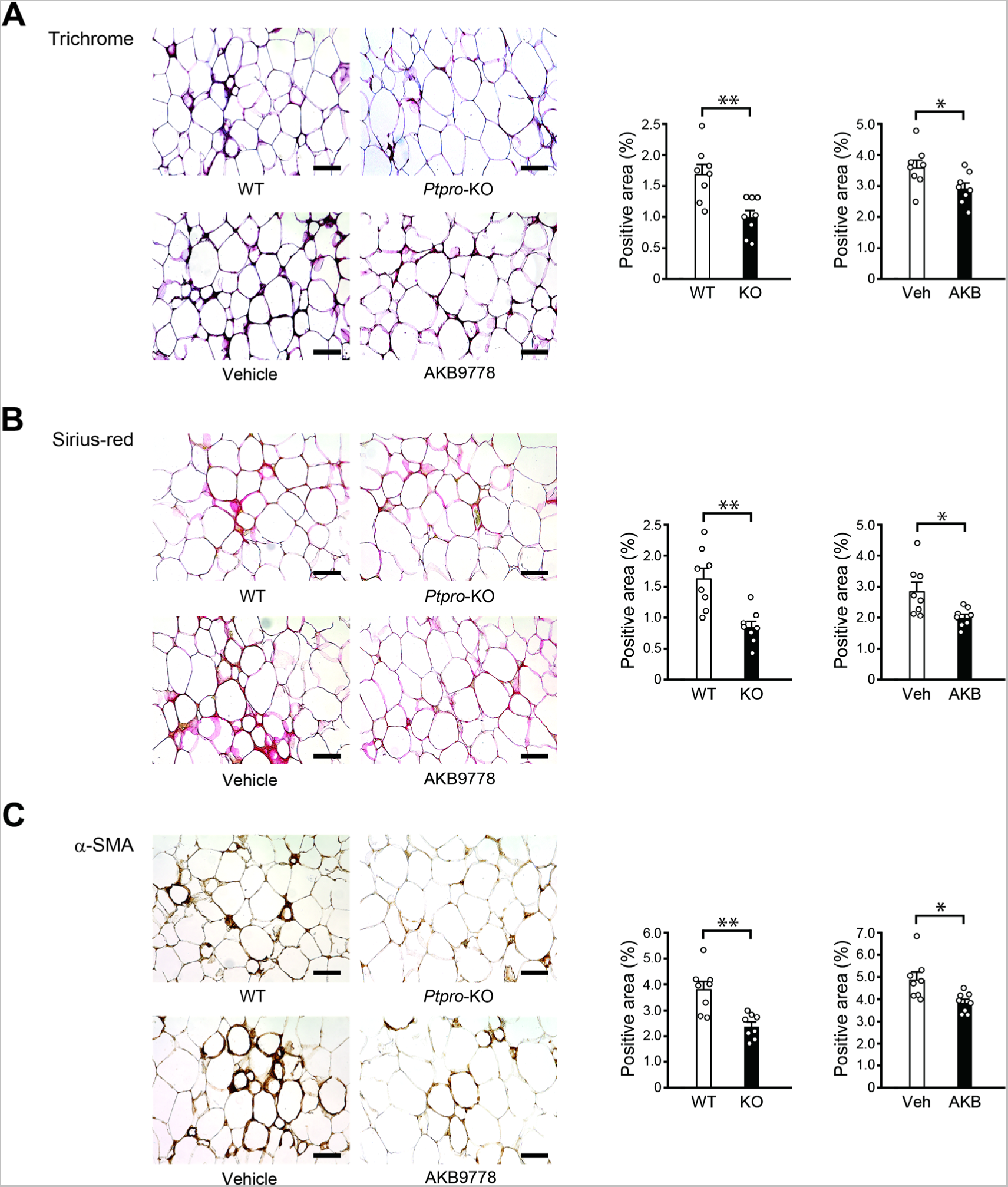
Reduced adipose tissue fibrosis in *Ptpro*-KO mice and AKB9778-treated *ob*/*ob* mice. (A) Masson’s trichrome staining of the epididymal adipose tissue of WT and *Ptpro*-KO mice fed HFHSD (upper) and *ob*/*ob* mice treated with vehicle or AKB9778 for 4 weeks (lower). Bars: 100 μm. The right graphs show quantitative analyses of positive areas (n = 8 each). (B) Picro-Sirius red staining of the epididymal adipose tissue of WT and *Ptpro*-KO mice fed HFHSD at 24 weeks of age (upper) and ND-fed *ob*/*ob* mice treated with vehicle or AKB9778 for 4 weeks (lower). Bars: 100 μm. The right graphs show quantitative analyses of positive areas (n = 8 each). (C) Anti-α-SMA immunostaining of the epididymal adipose tissue of WT and *Ptpro*-KO mice fed HFHSD (upper) and *ob*/*ob* mice treated with vehicle or AKB9778 (lower). Bars: 100 μm. The right graphs show quantitative analyses of positive areas (n = 8 each). Values are means ± SEM (A-C). *P* values are based on the unpaired Student’s *t*-test (A-C). \**P* < 0.05, \*\**P* < 0.01. **Source data 1.** Masson’s trichrome staining of the epididymal adipose tissue of WT and *Ptpro*-KO mice fed HFHSD and *ob*/*ob* mice treated with vehicle or AKB9778 (Figure 7A). **Source data 2.** Picro-Sirius red staining of the epididymal adipose tissue of WT and *Ptpro*-KO mice fed HFHSD and *ob*/*ob* mice treated with vehicle or AKB9778 (Figure 7B). **Source data 3.** Anti-α-SMA immunostaining of the epididymal adipose tissue of WT and *Ptpro*-KO mice fed HFHSD and *ob*/*ob* mice treated with vehicle or AKB9778 (Figure 7C).

### PTPRO is involved in the control of lipid accumulation in adipose tissue

We next examined the effect of PTPRO inhibition on adipocytes using the *in vitro* culture system of 3T3-L1 adipose cells. The sizes of fat droplets visibly increased in cells during adipocyte maturation with the AKB9778 treatment for 14 days (Figure 8A, upper). Quantitative analyses of the droplet diameter revealed that the number of small lipid droplets (2-4 μm in diameter) decreased (AKB9778; 30 ± 3: control; 74.6 ± 5.6 per 10^4^ µm^2^: *P* = 0.0004), whereas the number of large droplets (> 6 μm) increased (AKB9778; 30.4 ± 1: control; 11.2 ± 1.7 per 10^4^ µm^2^: *P* = 0.0002) with the AKB9778 treatment (Figure 8A, upper right). The average diameter of lipid droplets was 5.2 ± 0.1 μm in AKB9778-treated cells and 4.1 ± 0.2 μm in control cells (*P* = 0.0007) (Figure 8A, lower left). These results suggested that the AKB9778 treatment promoted the growth of lipid droplets in 3T3-L1 cells. The overall lipid content in a cell significantly increased in AKB9778-treated 3T3-L1 cells (*P* = 0.0125) (Figure 8A, lower right).

**Figure 8.**
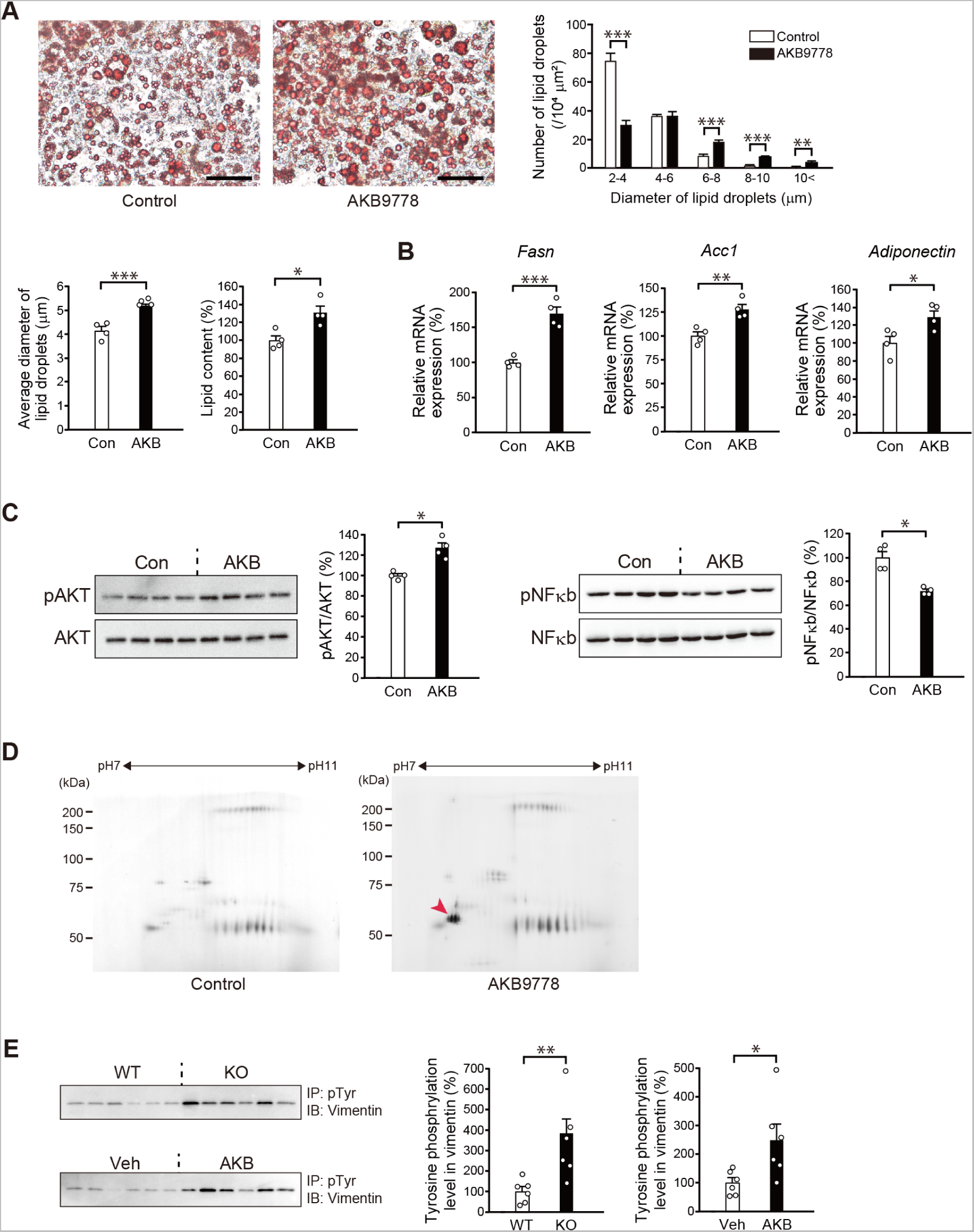
Reproduction of the AKB9778 effect in 3T3-L1 adipocytes *in vitro* and increase in the Tyr-phosphorylation of vimentin by the inhibition of PTPRO. (A) Oil red O staining of 3T3-L1 adipocytes treated with or without AKB9778 for 14 days in culture at the confluent state. Bars: 50 μm. The upper right graph shows the distribution of lipid droplets of the indicated sizes in 10^4^ μm^2^, and the lower left graph shows the averages of lipid droplet diameters (n = 4 each). The lower right graph shows lipid contents quantified by Oil Red O absorbance (n = 4 each). The result is given as a relative value to the control as a percentage. (B) mRNA expression levels of *Fasn*, *Acc1*, and *adiponectin* in 3T3-L1 adipocytes treated with or without AKB9778 for 14 days. n = 4 each. (C) Western blot analyses of pAKT and total AKT (left) and pNFκb and total NFκb in 3T3-L1 adipocytes treated with or without AKB9778 for 14 days. The graphs show quantitative analyses (n = 4 each). (D) Two-dimensional gel electrophoresis of Tyr-phosphorylated proteins prepared from 3T3-L1 adipocytes treated with or without AKB9778. Proteins are visualized by silver staining. The red arrowhead indicates Tyr-phosphorylated vimentin. (E) Western blot analyses of the phosphorylation of Tyr in vimentin in epididymal fat lysates prepared from WT and *Ptpro*-KO (KO) mice fed HFHSD at 24 weeks of age (upper). The phosphorylation of vimentin proteins was examined in immunoprecipitates with an anti-pTyr antibody with an anti-vimentin antibody. Western blot analyses of the phosphorylation of Tyr in vimentin in epididymal fat lysates prepared from ND-fed *ob*/*ob* mice treated with vehicle or AKB9778 for 4 weeks (lower). The right graphs show summaries of quantitative analyses (n = 6 each). Values are means ± SEM. *P* values are based on the unpaired Student’s *t*-test (A-E). \**P* < 0.05, \*\**P* < 0.01, \*\*\**P* < 0.001. **Source data 1.** Oil red O staining of 3T3-L1 adipocytes treated with or without AKB9778 for 14 days (Figure 8A). **Source data 2.** mRNA expression levels of *Fasn*, *Acc1*, and *adiponectin* in 3T3-L1 adipocytes treated with or without AKB9778 for 14 days (Figure 8B). **Source data 3.** Western blot analyses of pAKT and total AKT and pNFκb and total NFκb in 3T3-L1 adipocytes treated with or without AKB9778 for 14 days (Figure 8C). **Source data 4.** Western blot analyses of the phosphorylation of Tyr in vimentin in epididymal fat lysates prepared from WT and *Ptpro*-KO mice fed HFHSD and ob/ob mice treated with vehicle or AKB9778 (Figure 8E).

Consistent with this result, the expression levels of *Fasn* and *Acc1*, and *Adiponectin* were significantly higher in AKB9778-treated 3T3-L1 adipocytes than in control cells (*P* = 0.0005, *P* = 0.0054, and *P* = 0.0297, respectively) (Figure 8B), as observed in the adipose tissue of *Ptpro*-KO mice and AKB9778-treated *ob*/*ob* mice (Figure 3E, F). Furthermore, the activity of AKT was enhanced by the AKB9778 treatment in 3T3-L1 cells (*P* = 0.0024) (Figure 8C, left), whereas that of NFκb was suppressed (*P* = 0.0022) (Figure 8C, right). Collectively, the results obtained *in vivo* on the adipose tissue of *Ptpro*-KO mice and AKB9778-treated *ob*/*ob* mice were faithfully recapitulated in 3T3-L1 adipose cells treated with AKB9778 *in vitro*.

### PTPRO regulates Tyr phosphorylation levels in vimentin

To elucidate the molecular mechanism involving PTPRO in adipose tissue expansion, we examined differences in the phosphorylation of tyrosine (Tyr) in cellular proteins in 3T3-L1 adipose cells upon the AKB9778 treatment. By using the two-dimensional electrophoresis of Tyr-phosphorylated proteins, we found that a protein spot with ~53 kDa was markedly increased by the AKB9778 treatment (Figure 8D). This protein was identified as vimentin by mass spectrometry. Vimentin is a widely expressed intermediate filament protein that is reportedly associated with lipid droplets in adipocytes in adipose tissue (Franke et al., 1987; Olofsson et al., 2008). We also confirmed the phosphorylation of Tyr in vimentin in mouse adipose tissue. Tyr phosphorylation levels in vimentin were markedly higher in *Ptpro*-KO mice and AKB9778-treated *ob*/*ob* mice than in control mice (Figure 8E).

Several Tyr residues have been suggested to be phosphorylated in vimentin (Figure 9A, left); predominantly at Tyr(53), Tyr(61), and Tyr(117) (Cao et al., 2007; Yang et al., 2019). To identify the substrate site for PTPRO, we performed *in vitro* phosphatase assays using synthetic phosphopeptides corresponding to these Tyr residues in the mouse vimentin sequence (Figure 9A, middle). The phosphopeptide derived from the mouse insulin receptor, IR1, which was previously shown to be dephosphorylated by PTPRO (Shintani et al., 2015), was used as a positive control for the PTPRO substrate. PTPRO preferentially dephosphorylated the peptide containing phosphorylated Tyr(117) (Figure 9A, right), suggesting that PTPRO controls the dynamics of vimentin filaments through the dephosphorylation of Tyr(117) in vimentin.

**Figure 9.**
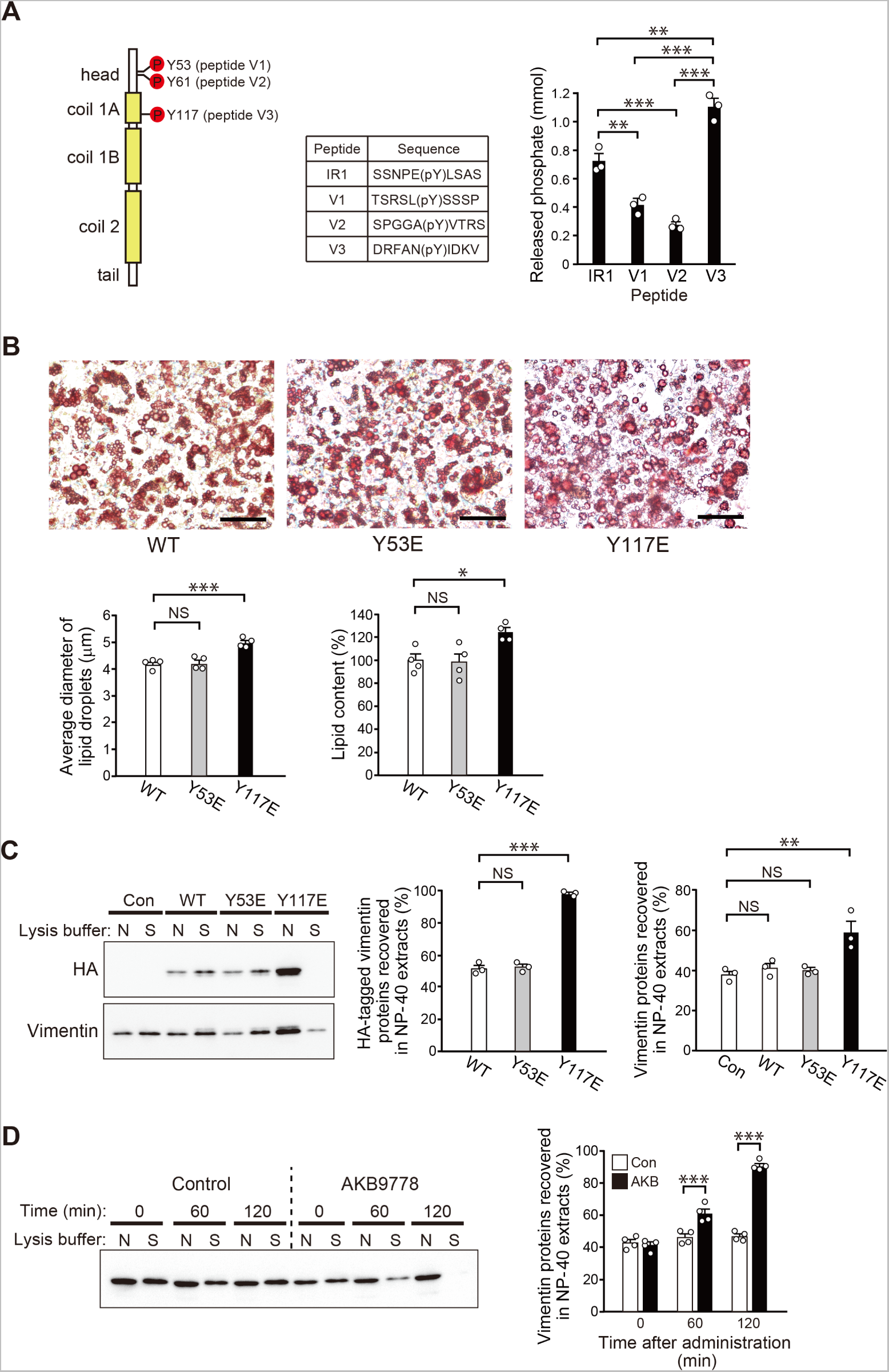
Functional regulation of vimentin by the phosphorylation of Tyr(117). (A) Dephosphorylation assays using synthetic phosphopeptides. Schematic representation of possible Tyr phosphorylation sites in mouse vimentin (left; the N terminus is the top). List of corresponding phosphopeptides (middle). Summary of phosphatase assays (right). Synthesized phosphopeptides were incubated with GST-PTPRO proteins, and the amount of phosphate released was measured (n = 3). Values are means ± SEM. *P* values are based on a one-way ANOVA followed by Tukey’s test. (B) Oil red O staining of 3T3-L1 adipocytes expressing the WT, Y53E mutant, or Y117E mutant of vimentin tagged with the HA epitope for 14 days at the confluent state. Bars: 50 μm. The lower left graph shows the average lipid droplet diameter (n = 4 each). The lower right graph shows lipid contents quantified by Oil Red O absorbance (n = 4 each). The result is given as a relative value to the control as a percentage. (C) Western blot analyses of vimentin by different extraction methods with the NP-40 lysis buffer (N) or SDS lysis buffer (S). Samples were prepared from 3T3-L1 adipocytes expressing the WT, Y53E mutant, or Y117E mutant of vimentin. Control cells were transfected with the empty vector. Exogeneous and total vimentin proteins were detected with an anti-HA antibody (upper left) and anti-vimentin antibody (lower left), respectively. The middle and right graphs show the results of quantitative analyses of HA-tagged vimentin and total vimentin recovered in NP-40 extracts (n = 3 each). (D) Western blot analyses of vimentin recovered in extracts with the NP-40 lysis buffer (N) or SDS lysis buffer (S) prepared from 3T3-L1 adipocytes treated with or without AKB9778. Cells were extracted at the indicated time points. The right graphs show summaries of quantitative analyses (n = 4 each). Values are means ± SEM. *P* values are based on a one-way ANOVA followed by Tukey’s test (A-D). NS, not significant; \**P* < 0.05, \*\**P* < 0.01, \*\*\**P* < 0.001. **Source data 1.** Dephosphorylation assays using synthetic phosphopeptides (Figure 9A). **Source data 2.** Oil red O staining of 3T3-L1 adipocytes expressing the WT, Y53E mutant, or Y117E mutant of vimentin (Figure 9B). **Source data 3.** Western blot analyses of vimentin by different extraction methods with the NP-40 lysis buffer or SDS lysis buffer (Figure 9C). **Source data 4.** Western blot analyses of vimentin recovered in extracts with the NP-40 lysis buffer or SDS lysis buffer prepared from 3T3-L1 adipocytes treated with or without AKB9778 (Figure 9D). **Figure supplement 1.** Forced expression of mutant proteins of vimentin in 3T3-L1 adipocytes. **Figure supplement 2.** Western blot analyses of vimentin recovered in extracts with the NP-40 lysis buffer (N) or SDS lysis buffer (S).

### Tyr(117) phosphorylation in vimentin stimulates the growth of lipid droplets in adipocytes

To clarify the role of the phosphorylation of Tyr(117) in vimentin, we established 3T3-L1 cells expressing the HA-tagged Y117E mutant, in which the Tyr residue was converted to glutamate to mimic phosphorylation. 3T3-L1 cells expressing the HA-tagged WT or Y53E mutant were also prepared as a control. After the differentiation of 3T3-L1 adipocytes in the culture, exogenous vimentin expression was induced for 14 days by the doxycycline treatment and detected using the HA epitope at the C terminus (Figure 9-figure supplement 1A). Y117E-expressing cells contained larger lipid droplets than WT-expressing cells or Y53E mutant-expressing cells used as negative controls (Figure 9B, upper); the average diameters of lipid droplets were 5.0 ± 0.1, 4.2 ± 0.1, and 4.2 ± 0.1 μm in Y117E, Y53E, and WT-expressing 3T3-L1 cells, respectively (*P* = 0.0007 compared with WT) (Figure 9B, lower left). Consistently, the overall lipid content was significantly higher in Y117E-expressing cells (*P* = 0.0349) (Figure 9B, lower right). These results suggested that the phosphorylation of Tyr(117) in vimentin plays a pivotal role in the growth of lipid droplets.

Y117E mutant proteins showed a spotty distribution in 3T3-L1 adipocytes, whereas WT and Y53E mutant proteins had a fibrous distribution (Figure 9-figure supplement 1B). When 3T3-L1 adipocytes were extracted with lysis buffers containing NP-40 or SDS as the detergent, endogenous vimentin proteins were recovered in both fractions, as well as WT and Y53E mutant proteins. However, the majority of Y117E mutant proteins were recovered in the NP-40 fraction (Figure 9C), suggesting that the Y117E mutant did not form intermediate filaments. The distribution ratio in the NP-40 fraction of total vimentin was also increased by Y117E mutant expression in 3T3-L1 adipocytes (Figure 9C, lower left and right), indicating that the expression of the Y117E mutant exerted negative effects on the filament formation of vimentin.

The percentage of vimentin recovered in the NP-40 fraction increased with time after the AKB9778 treatment on 3T3-L1 cells, and the majority of vimentin was recovered in this fraction at 120 min (Figure 9D). When adipose tissues were analyzed, the percentage of vimentin recovered in the NP-40 fraction was expectedly higher in *Ptpro*-KO mice and AKB9778-treated *ob*/*ob* mice than in control mice (Figure 9-figure supplement 2). These results suggest that PTPRO activity promotes the formation of vimentin filaments by reducing the phosphorylation level at Tyr(114).

### The *Ptpro* deficiency and AKB9778 treatment improve lipid and glucose homeostasis

To estimate systemic conditions of HFHSD-fed *Ptpro*-KO mice and AKB9778-treated *ob*/*ob* mice, we examined blood lipid concentrations in the sera. Serum from *Ptpro*-KO mice showed significantly lower levels of TG, non-esterified fatty acids (NEFA), and total cholesterol (T-Cho) than the serum of WT mice (Figure 10A, upper). *Ob/ob* mice showed markedly high levels of TG, NEFA, and T-Cho; however, AKB9778-treated *ob*/*ob* mice showed significantly reduced lipid levels of all types (Figure 10A, lower). These results indicate that the *Ptpro* deficiency and AKB9778 treatment improved lipid homeostasis.

**Figure 10.**
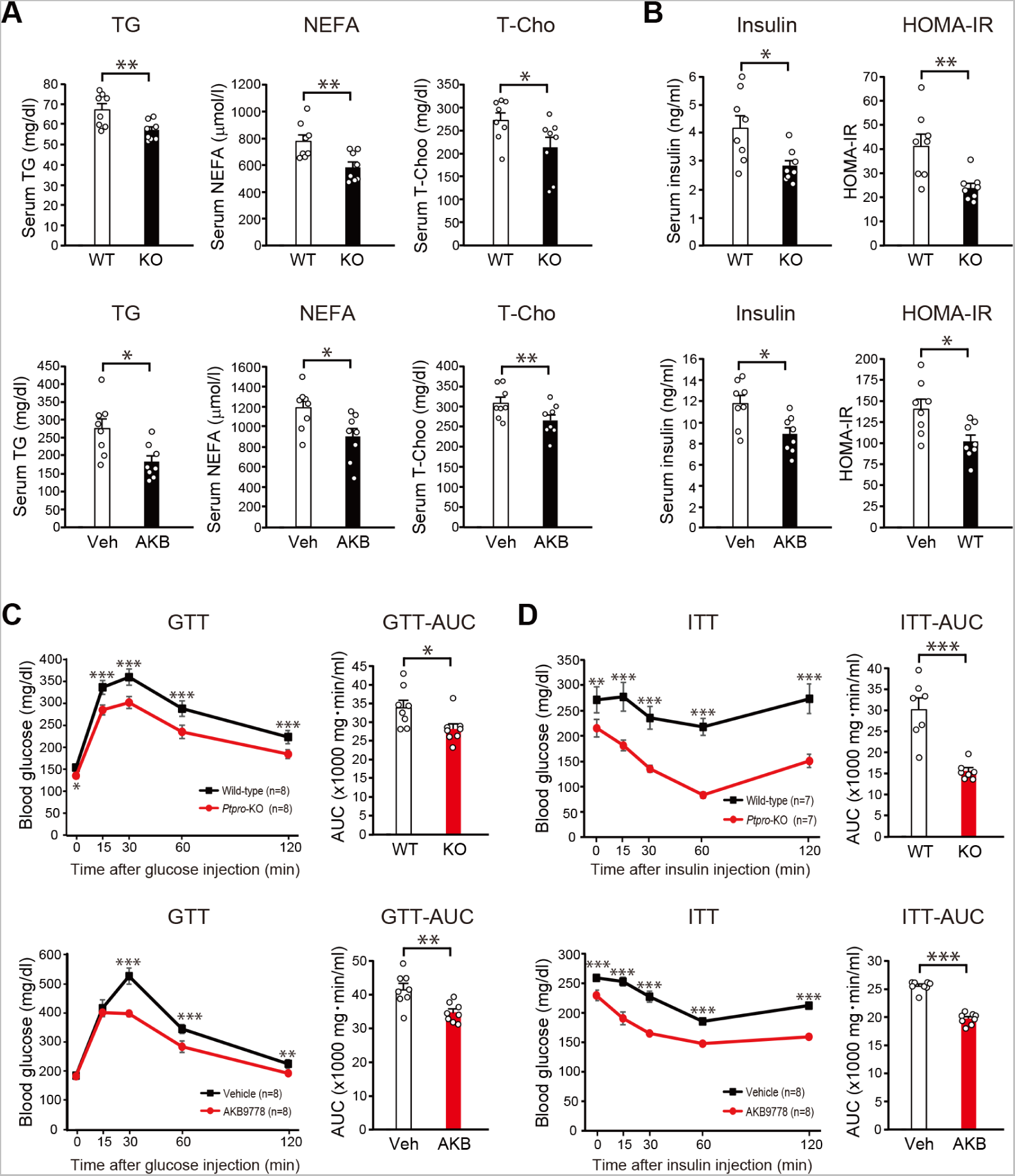
Improved lipid and glucose homeostasis in *Ptpro*-KO mice and AKB9778-treated *ob*/*ob* mice. (A) Serum levels of triglycerides (TG), non-esterified fatty acids (NEFA), and total cholesterol (T-Cho) in WT and *Ptpro*-KO (KO) mice at 24 weeks of age fed HFHSD (upper), or in *ob*/*ob* mice fed ND after the treatment with vehicle (Veh) or AKB9778 (AKB) for 4 weeks from 16 weeks of age (lower). n = 8 each. (B) Serum insulin levels and HOMA-IR levels in WT and *Ptpro*-KO mice fed HFHSD (upper), or in *ob*/*ob* mice treated for 4 weeks with vehicle or AKB9778 (lower). (C) The glucose tolerance test (GTT) in WT and *Ptpro*-KO mice fed HFHSD (upper), or in *ob*/*ob* mice treated with vehicle or AKB9778 (lower). (D) The insulin tolerance test (ITT) in WT and *Ptpro*-KO mice fed HFHSD (upper), or in *ob*/*ob* mice treated with vehicle or AKB9778 (lower). Values are means ± SEM. The *P* values of GTT and ITT curves were analyzed by a two-way ANOVA followed by Tukey’s test, and the *P* values of other data were based on the unpaired Student’s *t*-test. \**P* < 0.05, \*\**P* < 0.01, \*\*\**P* < 0.001. **Source data 1.** Serum levels of triglycerides, non-esterified fatty acids, and total cholesterol in WT and *Ptpro*-KO mice, or in *ob*/*ob* mice treated with vehicle or AKB9778 (Figure 10A). **Source data 2.** Serum insulin levels and HOMA-IR levels in WT and *Ptpro*-KO mice, or in *ob*/*ob* mice treated with vehicle or AKB9778 (Figure 10B). **Source data 3.** The glucose tolerance test in WT and *Ptpro*-KO mice, or in *ob*/*ob* mice treated with vehicle or AKB9778 (Figure 10C). **Source data 4.** The insulin tolerance test in WT and *Ptpro*-KO mice, or in *ob*/*ob* mice treated with vehicle or AKB9778 (Figure 10D).

Circulating insulin regulates many aspects of homeostasis, including lipid and carbohydrate metabolism. Under obese conditions, insulin levels in fasting blood were significantly lower in *Ptpro*-KO mice than in WT mice (*Ptpro*-KO; 2.79 ± 0.2 ng/ml: WT; 4.15 ± 0.44 ng/ml: *P* = 0.0136) (Figure 10B, upper left). Furthermore, homeostasis model assessment of insulin resistance (HOMA-IR) levels, a marker of insulin resistance, were markedly lower in *Ptpro*-KO mice than in WT mice (*P* = 0.0049) (Figure 10B, upper right). AKB9778-treated *ob*/*ob* mice also showed markedly lower insulin levels (AKB9778; 8.89 ± 0.61 ng/ml: Vehicle; 11.79 ± 0.77 ng/ml: *P* = 0.0103) and HOMA-IR levels than those in vehicle mice (*P* = 0.0114) (Figure 10B, lower). These results indicated significant improvements in insulin resistance in *Ptpro*-KO mice and AKB9778-treated *ob*/*ob* mice, although both of them were obese.

To assess the blood glucose control and insulin sensitivity, we conducted a glucose tolerance test (GTT) and insulin tolerance test (ITT). Increases in blood glucose levels induced by the intraperitoneal administration of glucose (GTT) were more gradual in *Ptpro*-KO mice than in WT mice (Figure 10C, upper left); the area under the curve (AUC) of GTT was significantly smaller in *Ptpro*-KO mice (*P* = 0.0256) (Figure 10C, upper right). Blood glucose increases following the intraperitoneal glucose challenge (GTT) were consistently smaller in AKB9778-treated *ob*/*ob* mice than in vehicle mice (Figure 10C, lower). In addition, a more pronounced decrease in blood glucose levels after the administration of insulin (ITT) was observed in *Ptpro*-KO mice than in WT mice (Figure 10D, upper left); at 60 min, blood glucose levels decreased by ~70% in *Ptpro*-KO mice and by ~20% in WT mice (*P* < 0.0001). The AUC of ITT was accordingly smaller in *Ptpro*-KO mice than in WT mice (Figure 10D, upper right). The decreases induced in blood glucose levels by the administration of insulin (ITT) were consistently pronounced in AKB9778-treated mice (*P* < 0.0001) (Figure 10D, lower). Therefore, glucose homeostasis and insulin sensitivity were clearly ameliorated in obese *Ptpro*-KO mice and AKB9778-treated *ob*/*ob* mice.

## DISCUSSION

Diet-induced obesity is a key risk factor for many chronic illnesses, including liver steatosis, cardiovascular diseases, and T2DM (Goossens, 2017; Longo et al., 2019). In the adipose tissue of obese animals, the capacity for fat synthesis is generally reduced, whereas that for fat degradation is increased, which leads to the release of free fatty acids from adipose tissue. As a result, the accumulation of ectopic fat occurs in other tissues, such as the liver, which is the pathogeny of lipotoxicity (Goossens, 2017; Kahn et al., 2019). Saturated fatty acids released from the adipocytes of obese animals are recognized by TLR4, an innate immune receptor expressed on macrophages, which leads to the activation of NFκb and the induction of inflammation (Lee et al., 2001; Shi et al., 2006; Suganami et al., 2007). Inflammation and insulin resistance are also linked and aggravate the progression of metabolic disorders (Goossens, 2017; Longo et al., 2019). Pro-inflammatory cytokines, such as TNFα, induce insulin resistance and decrease the cellular utilization of glucose (Hotamisligil et al., 1993). In the present study, we analyzed HFHSD-fed *Ptpro*-KO mice and ND-fed *ob*/*ob* mice treated with AKB9778, an inhibitor of PTPRO, and revealed that PTPRO plays a pivotal role in the control of metabolism and inflammation in the liver and adipose tissue, as well as the systemic dysfunction associated with obesity.

In PTPRO-deficient mice, inflammation in adipose tissue (Figure 4) and the liver (Figure 2) was markedly low and the metabolic state remained healthy (Figures 2, 3), despite the significant expansion of adipose tissue (Figure 1). In more details, the accumulation of ectopic fat in the liver was markedly lower than in control mice (Figure 1) and the expression of pro-inflammatory factors was significantly suppressed (Figures 2, 4), while on the other hand, lipid and glucose homeostasis (Figures 1, 10) and insulin sensitivity (Figure 10) were almost normal or only slightly impaired. More importantly, the administration of AKB9778, the inhibitor of PTPRO, to hyper obese *ob*/*ob* mice induced the redistribution of lipids from the ectopic fat storage to adipose tissue (Figures 1, 3) by the induction of lipogenic genes (Figure 3) and promotion of the growth of lipid droplets in adipocytes (Figure 8), which resultantly improved systemic inflammation and insulin sensitivity (Figures 4, 10).

PTPRO activity is assumed to be involved in the metabolic function of tissues throughout the body by the dephosphorylation of multiple substrate molecules, and accordingly, its mode of action may be complex. Several PTKs, including the insulin receptor, FGF receptor, and EGF receptor, and Abl have been identified as substrates for PTPRO, at least at the molecular level (Chen et al., 2006; Sakuraba et al., 2013; Shintani et al., 2006; Shintani et al., 2015; Shintani et al., 2017; Yu et al., 2018). Therefore, increases in insulin sensitivity by AKB9778 may be primarily attributed to the direct inhibition of PTPRO and PTPRJ in the insulin target tissues, such as skeletal muscle and liver, because they negatively regulate insulin receptors through dephosphorylation *in vivo* (Shintani et al., 2015). The present study demonstrated that AKT was significantly activated in the adipose tissue of obese *Ptpro*-KO and AKB9778-treated *ob*/*ob* mice (Figure 3G) and AKB9778-treated 3T3-L1 adipocytes (Figure. 8C). The PI3K/AKT signaling pathway is downstream of the insulin receptor and has been shown to promote glucose uptake into cells and lipid biosynthesis and also inhibit lipolysis (Huang et al., 2018). The expression of lipogenic enzymes was consistently enhanced, while that of enzymes for lipolysis was suppressed in the adipose tissue of obese *Ptpro*-KO and AKB9778-treated *ob*/*ob* mice (Figure 3). Therefore, enhanced PI3K/AKT signaling in *Ptpro*-KO and AKB9778-treated *ob*/*ob* mice may explain adipose tissue expansion and reductions in free FA levels in blood.

NFκb is a transcription factor that is involved in the regulation of a wide range of genes related to inflammation. Obesity is associated with the activation of the NFκb inflammatory pathway (Kahn et al, 2019; Napetschnig; 2013). PTPRO was preferentially expressed in adipocytes and macrophages in adipose tissue (Figure 1-figure supplement 3). Importantly, PTPRO has been suggested to play a role in the activation of NFκb in macrophages; it is involved in the pathogenesis of ulcerative colitis by activating the TLR4/NFκb signaling pathway in macrophages (Zhao et al., 2020). PTPROt, the truncated form of PTPRO, in liver macrophages is also reportedly involved in the exacerbation of steatohepatitis via the activation of NFκb (Jin et al., 2020); however, the molecular mechanisms by which PTPROt aggravates inflammation remain largely unknown. In contrast, the fundamental role of PTPRO in adipocytes has not yet been studied. We herein demonstrated that the activation of NFκb was suppressed in the adipose tissue of obese *Ptpro*-KO mice and AKB9778-treated *ob*/*ob* mice (Figure 5E). Moreover, NFκb activity was reduced in AKB9778-treated 3T3-L1 adipocytes (Figure 8C). The lack or inhibition of PTPRO activity may similarly suppress the activation of NFκb in adipocytes as well as in macrophages.

Inflammation was suppressed in the adipose tissue and liver of obese *Ptpro*-KO mice and AKB9778-treated *ob*/*ob* mice, and the infiltration of macrophages into adipose tissue was also attenuated (Figures 2, 4, 5). The expression levels of pro-inflammatory factors, such as *Tnfa*, *IL6*, and *Mcp1*, were accordingly suppressed in the liver and adipose tissue in these mice (Figures 2, 4). *Ptpro*-KO mice and AKB9778-treated mice showed low levels of TNFα and IL6 also in blood (Figure 4). The pro-inflammatory cytokines released from macrophages exert pathogenic effects on other tissues. A previous study demonstrated that TNFα induced insulin resistance and decreased cellular glucose utilization (Hotamisligil et al., 1993). When inflammatory responses are initiated in adipose tissue, hypertrophied adipocytes also release pro-inflammatory adipokines and further exacerbate the inflammatory response (Kahn et al., 2019; Napetschnig and Wu, 2013; Suganami et al., 2020). The formation of crown-like structures was suppressed in *Ptpro*-KO mice and AKB9778-treated *ob*/*ob* mice (Figure 5), indicating that cell death in adipocytes was also reduced. PTPRO expressed in adipocytes and macrophages may be cooperatively involved in the systemic regulation of inflammatory responses.

PTP1B is widely expressed in multiple tissues, including adipose tissue and the liver (Bourdeau et al., 2005). PTP1B is known to suppress insulin signaling by direct dephosphorylation of insulin receptor (Seely et al., 1996). Mice lacking PTP1B show increased insulin signaling and sensitivity (Elchebly et al., 1999). The improvements in insulin sensitivity in obese *Ptpro*-KO mice and AKB9778-treated *ob*/*ob* mice may be attributable in part to the suppression of PTP1B expression. In addition, *Ptp1b*-KO mice are protected from diet-induced obesity owing to enhanced leptin action in the hypothalamus and have reduced adiposity (Cheng et al., 2002; Zabolotny et al., 2002). PTP1B expression is increased in adipose tissue and the hypothalamus by inflammation induced by obesity (Nieto-Vazquez et al., 2008; Zabolotny et al., 2008). PTP1B is reportedly upregulated via TNFα (Ahmal et al., 1997) and high-fat diet feeding of mice leads to concomitant increase in both TNFa and PTP1B levels (Compa et al., 2002). We also observed that HFHSD-feeding increased *Ptp1b* expression in adipose tissue of WT mice (Figure 4A). However, in obese *Ptpro*-KO mice and AKB9778-treated *ob*/*ob* mice, *Ptp1b* expression was significantly suppressed, probably due to a reduced inflammation (Figure 4A, B). Therefore, the expression of *Ptp1b* induced by inflammation in obesity is thought to be downstream and under the control of PTPRO in a broad sense.

The ECM is considered to be an important factor in maintaining the structural integrity of adipocytes under normal conditions (O’Connor et al., 2003; Strieder-Barboza et al., 2020). However, inflammation associated with obesity often leads to remodeling of the ECM with fibrosis. In the adipose tissue of obese animals, fibrosis has been shown to impair the physiological functions of adipose tissue as a nutrient storage organ and induce insulin resistance (Sun, et al., 2013; Marcelin et al., 2019). Therefore, the inhibition of fibrosis in adipose tissue is expected to ameliorate the metabolic abnormalities associated with obesity. When the collagen VI gene, a major component of the ECM of adipose tissue, was additionally deficient in *ob*/*ob* mice, the expansion of individual adipocytes was not constrained, and the resultant expansion of adipose tissue with enlarged adipocytes was associated with improved systemic insulin sensitivity (Kahn et al., 2009). Similarly, when lysyl oxidase, an enzyme involved in collagen cross-linking, was inhibited in obese mice, inflammation in adipose tissue was reduced and glucose homeostasis was improved (Halberg et al., 2009). Notably, expression levels of ECM-related genes and fibrosis in adipose tissue was markedly suppressed in *Ptpro*-KO and AKB9778-treated *ob*/*ob* mice (Figures 6 and 7). Mild fibrosis may also be one of the mechanisms underlying the enlargement of adipocytes associated with improved metabolism in these mice.

The present study revealed that the phosphorylation of vimentin was elevated in the adipose tissue of obese *Ptpro*-KO and AKB9778-treated *ob*/*ob* mice (Figure 9). Vimentin belongs to intermediate filaments and assembles into a network of filaments that spans the cytoplasm (Lowery et al., 2015). Vimentin has been reported to bind to cellular organelles, including the Golgi, mitochondrion, endoplasmic reticulum, and nucleus, and regulates and stabilizes the organization of these organelles (Guo et al., 2013). A previous study reported that the phosphorylation of Tyr(117) in vimentin by Src regulated the dynamics and organization of vimentin filaments during cell migration (Yang et al., 2019). In mature adipocytes, lipid droplets are covered by cages of vimentin intermediate filament (VIF) (Heid et al., 2014). We demonstrated that PTPRO regulated the phosphorylation level of Tyr(117) in vimentin (Figure 9), and the phosphorylated form of vimentin led to the disruption of vimentin filaments (Figure 9). Larger lipid droplets formed in AKB9778-treated 3T3-L1 cells than in control cells (Figure 8), suggesting that the phosphorylation of Tyr(117) in vimentin loosens VIF cages and promotes the growth of lipid droplets in adipocytes.

We herein revealed by *in vivo* assay that AKB9778 exhibited high inhibitory activity against R3 members (PTPRO, PTPRJ, and PTPRB), particularly PTPRO (Figure 1-figure supplement 4). The primary target of AKB9778 *in vivo* expectedly appears to be PTPRO because the majority of phenotypes observed in *Ptpro*-KO mice were reproduced in AKB9778-treated *ob*/*ob* mice. The overeating behavior against HFHSD observed in *Ptpro*-KO mice was not expectedly observed by AKB9778-treated *ob*/*ob* mice (Figure 1A and Shintani et al., 2015). It is noteworthy that *Ptprj*-KO mice show a lean phenotype, the opposite phenotype to *Ptpro*-KO mice, because PTPRJ directly suppresses leptin signaling (Shintani et al., 2017). Moreover, AKB9778 cannot cross the blood-brain barrier due to its highly hydrophilic structure (Stefani et al., 2016), and thus, AKB9778 may not affect the activity of PTPRO or PTPRJ in the hypothalamic feeding center in the brain. This may be the reason why the eating behavior of *ob*/*ob* mice was not affected following the treatment with AKB9778.

In summary, the present study demonstrated for the first time that a single drug targeting PTPRO improved ectopic fat accumulation in the liver and ameliorated the metabolic complications associated with obesity. Increases in the adipose tissue mass beyond normal limits, along with the larger size of adipocytes, were associated with the suppression of inflammation, as well as improvements in insulin resistance and glucose metabolism. This is consistent with the notion that metabolic activities and inflammation are highly related (Abreu-Vieira et al., 2015; Kim et al., 2008; Tanaka et al., 2014; Verschoor et al., 2021). We previously reported that PTPRO suppressed insulin signaling through the inactivation of insulin receptors (Shintani et al., 2015). In the present study, we showed *in vivo* and at the cellular level that PTPRO was involved in regulating the expression of lipogenic genes, the growth of lipid droplets in adipocytes, and the induction of inflammation at the tissue and systemic levels. Therefore, PTPRO may be a hub regulator for the development of chronic inflammation, insulin resistance, dyslipidemia, diabetes mellitus, and fatty liver associated with obesity.

## MATERIALS AND METHODS

### Experimental animals

All experiments with mice were performed according to protocols approved by the Institutional Animal Care and Use Committee of the National Institutes of Natural Sciences, Japan (approval numbers 18A031 and 19A037) and the Tokyo Institute of Technology (approval number D2019013), which are consistent with the ARRIVE guidelines. C57BL/6J and C57BL/6-Lepob/J (*ob*/*ob*) mice were purchased from Nippon Clea (Tokyo, Japan) and Charles River Laboratories Japan (Yokohama, Japan), respectively. *Ptpro*-KO mice were obtained from the RIKEN BioResource Center (catalog number 01235, RRID: IMSR_RBRC01235). *Ptpro*-KO mice were originally generated in the laboratory of one of the authors, T. Shirasawa, and backcrossed with C57BL/6J mice to obtain littermates for analyses. Mice were housed in a pathogen-free environment at a room temperature of 24 ± 1°C and humidity of 40-60% with a flickering cycle of 8:00-20:00. Co-habiting mice of both sexes were housed in plastic cages (cage size: 12 × 21 × 12.5 cm) filled with wood shavings and fed chow diet CA-1 (Nippon Clea, Tokyo, Japan) or HFHSD (F2HFHSD, Oriental Yeast Co., Ltd., Tokyo, Japan) consisting of protein (17.2% of energy intake), fat (54.5%), and carbohydrate (28.3%). All studies compared WT and *Ptpro*-KO littermate mice, or *ob*/*ob* mice treated with vehicle or AKB9778, which confers group allocation and randomisation in cage housing.

### Histological examination

After perfusion fixation with 10% formaldehyde, mouse tissues were removed and post-fixed in 10% formaldehyde at 4°C overnight. Tissues were then dehydrated in a graded ethanol series (50–100%) and embedded in paraffin. Six-micrometer-thick sections were deparaffinized and stained with hematoxylin and eosin, the Masson’s Trichrome Stain Kit (Scy Tek Laboratories, USA), and Picro-Sirius Red Stain Kit (Scy Tek Laboratories, USA), or immunostained with the antibodies shown in the Table 3.

**Table 3.** Antibodies used for immunohistochemistry and Western blot.

| Target protein | Source | Dilution for WB | Dilution for IHC |
| --- | --- | --- | --- |
| AKT (pan) | CST: #4691 | 1:2000 |  |
| Phospho-AKT (pan) | CST: #4060 | 1:1000 |  |
| F4/80 | Santa Cruz:<br>sc-52654 |  | 1:500 |
| NFkb | CST: #30656 | 1:1000 |  |
| Phospho-NFkb | CST: #4025 | 1:1000 |  |
| Phosphotyrosine | Merck: 05-321 | 1:1000 |  |
| Vimentin | CST: #5741 | 1:1000 | 1:500 |
| PTPRO | A gift from Prof.<br>T. Matozaki | 1:500 | 1:200 |
| Perilipin | CST: #9349 |  | 1:500 |
| CD11c | CST: #97585 |  | 1:500 |
| HA | CST: #3724 | 1:1000 | 1:500 |

Immunostaining was performed as previously described (Shintani et al., 2017). Images were acquired with a BZ-X810 fluorescence microscope (Keyence Corporation, Osaka, Japan). Stained areas were quantified in 4 non-overlapping fields randomly selected in each animal using BZ-H4C imaging analysis software (Keyence Corporation, Osaka, Japan), and the results obtained were expressed as a percentage of the positive area. The cell diameter of adipocytes was estimated using the same software for 100-150 adipocytes per mouse.

In Bodipy staining, after perfusion fixation with 10% formaldehyde, livers were removed and post-fixed in 10% formaldehyde at 4°C overnight. They were then infiltrated with a graded sucrose series to 18%, embedded in OCT compound (Miles Scientific, Naperville, IL), and sectioned at a thickness of 16 μm. Sections were incubated in Bodipy 493/503 (Invitrogen, USA) diluted to a concentration of 1 μg/ml in PBS at room temperature for 20 min. Sections were then washed twice with PBS and treated with Fluoromount (Diagnostic BioSystems, USA) for inclusion. Fluorescence images were acquired with a Keyence BZ-X810 fluorescence microscope.

### Analysis of the inhibitory activity of AKB9778 against PTPs

AKB9778 was synthesized by Sundia MediTech (Shanghai, China). The intracellular regions of PTPRO, PTPRJ, and PTPRB were prepared using the *Escherichia coli* expression system as previously described (Shintani et al., 2006) and the intracellular regions of PTPRC, PTPRD, and PTP-1B were prepared using the silkworm expression system as previously described (Fujikawa et al., 2017). The dephosphorylation reaction was initiated by adding DiFMUP (Thermo Fisher), an artificial substrate for phosphatases, to solutions of individual PTP proteins in Hepes buffer (50 mM Hepes, 100 mM NaCl, 2 mM EDTA, and 1 mM DTT, pH 7.0) containing various concentrations of AKB9778. The reaction was performed at 37°C for 30 min. The inhibitory activity of AKB9778 was estimated by measuring fluorescence values using the Corona multigrating microplate reader SH-9000Lab (Hitachi High-Tech Science, Tokyo, Japan) with excitation/emission wavelengths of 350/455 nm.

### *In vivo* administration of AKB9778

AKB9778 was dissolved at a concentration of 5 mg/ml in 10% HPCD (2-hydroxypropyl-β-cyclodextrin) (Nacalai Tesque, Kyoto, Japan) and intraperitoneally administered to *ob*/*ob* mice from 16 to 20 weeks of age at a dose of 10 mg/kg body weight daily between 8:00 and 9:00. In the vehicle group, only 10% HPCD was administered at the same dose.

### Treatment of 3T3-L1 cells with AKB9778 and analysis of fat accumulation

Mouse 3T3-L1 pre-adipocytes were purchased from the American Type Cell Collection (Manassas, VA, USA; ID: CL-173). They were grown using Dulbecco’s modified Eagle’s medium (DMEM) containing 10% fetal bovine serum (FBS) at 37°C under an atmosphere of 5% CO_2_. From the third day of postconfluence, 3T3-L1 cells were incubated with DMEM containing 10% FBS, 20 μg/ml insulin, 1 μM dexamethasone, and 0.5 mM isobutylmethylxanthine to induce differentiation for 2 days, and then with DMEM containing 10% FBS and 1 μg/ml insulin supplemented with or without 10 μM AKB9778 for an additional 14 days. AKB9778 was dissolved in dimethyl sulfoxide (DMSO), and the final DMSO concentration in the medium was 0.1%. In the control culture, only DMSO was added.

In the analysis of fat accumulation in 3T3-L1 adipocytes, cells were fixed with 10% buffered formalin. Cells were stained with 0.2% Oil-red O solution at room temperature for 10 min and washed with 60% isopropanol and PBS. Images were acquired with a BZ-X810 microscope (Keyence Corporation, Osaka, Japan), and the diameter of lipid droplets was measured in 10^4^ μm^2^ of 3 non-overlapping fields randomly selected from each experiment (n = 4) using BZ-H4C imaging analysis software (Keyence Corporation, Osaka, Japan). Lipid droplets larger than 2 μm in diameter were counted. To assess lipid quantities in cells, the dye was dissolved in isopropanol and absorbance at 510 nm was measured.

### Quantitative analysis of total lipids, TG, and cholesterol in the liver

Liver lipids were extracted according to the method of Folch et al. (Folch et al., 1957). Total lipids were assessed by dry weight. The lipid extract was dissolved in isopropanol and the amounts of TG and cholesterol were measured using the LabAssay Triglyceride (Fujifilm Wako Pure Chemical Corporation, Osaka, Japan) and Cholesterol (Fujifilm Wako Pure Chemical Corporation, Osaka, Japan) kits, respectively.

### Real-time quantitative PCR

Total RNA was extracted using TRIzol Reagent (Life Technologies, USA) according to the manufacturer’s instructions and reverse transcribed using the PrimeScript RT Reagent Kit with the gDNA Eraser (Takara, Shiga, Japan). To amplify cDNA, SYBR Premix Ex TaqII (Takara, Shiga, Japan) was used in the StepOnePlus Real-Time PCR thermocycler (Applied Biosystems, USA). PCR settings were as follows: after initial denaturation at 95°C for 30 s, amplification was performed at 95°C for 5 s and at 60°C for 30 s for 40 cycles. The quantification of gene expression was calculated relative to *Gapdh*. Primer sequences are shown in the following Table 4.

**Table 4.**
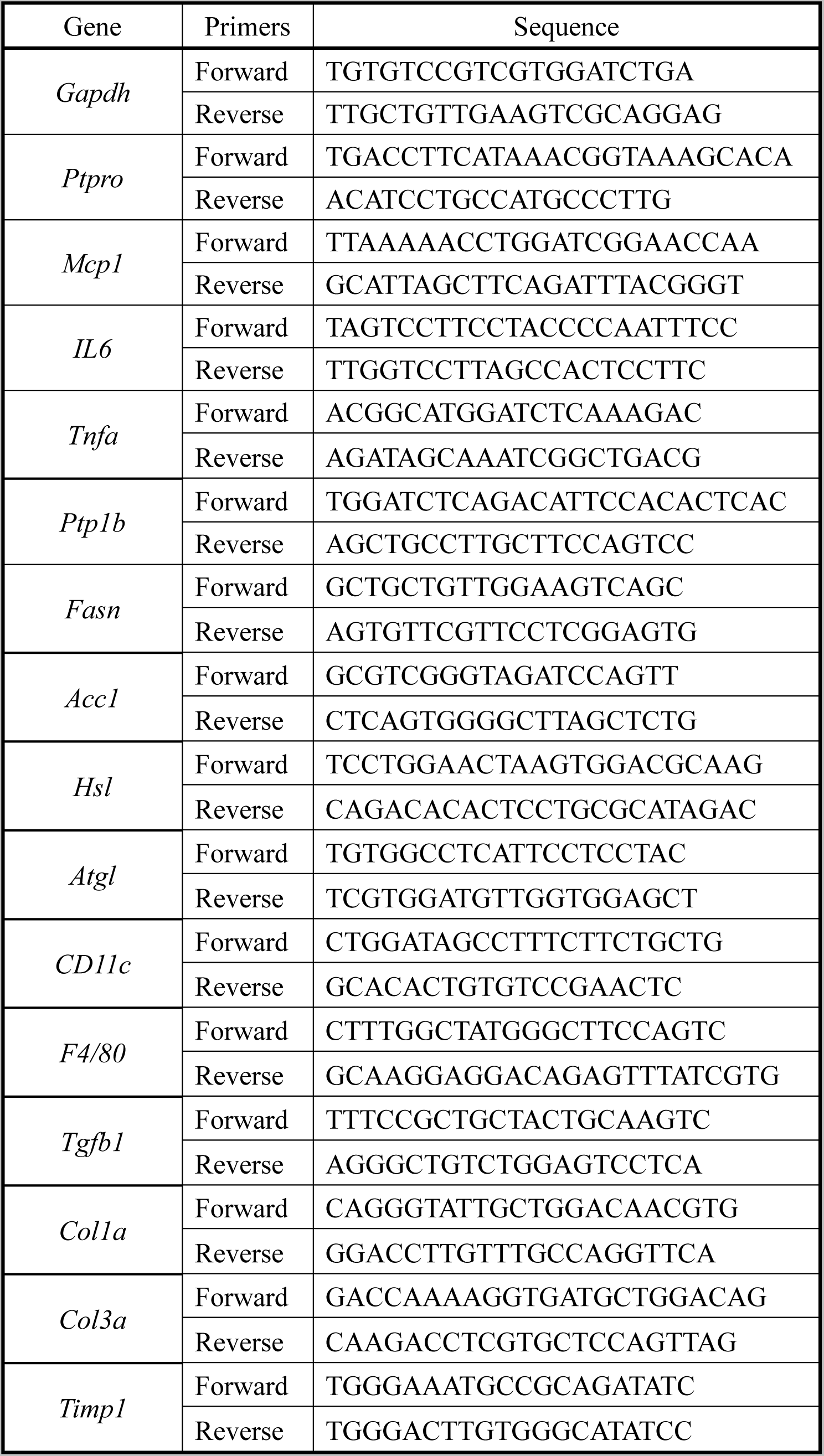
Sequences of quantitative PCR primers.

### Analysis of blood parameters

Blood was placed on ice for 30 minutes, and then centrifuged at 1200 × *g* for 20 minutes to obtain serum. TG, cholesterol, and NEFA concentrations in serum were measured using the Lab Assay Triglyceride, Cholesterol, and NEAA Kits (Fujifilm Wako Pure Chemical Corporation, Osaka, Japan), respectively. Serum ALT, AST, TNFα, IL6, adiponectin, insulin, and leptin concentrations were assayed using the Mouse ALT ELISA Kit (Abcam, USA), Mouse AST ELISA Kit (Abcam), Mouse TNF-alpha Quantikine ELISA Kit (R & D Systems, USA), Mouse IL-6 Quantikine ELISA Kit (R & D Systems, USA), Revis Adiponectin-Mouse/Rat ELISA Kit (Fujifilm Wako Pure Chemical Corporation, Osaka, Japan), Morinaga Insulin ELISA Kit (Morinaga Institute of Biological Science, Yokohama, Japan), and Morinaga Leptin ELISA Kit (Morinaga Institute of Biological Science, Yokohama, Japan), respectively. HOMA-IR values were calculated as follows: (fasting insulin level (µIU/ml) × fasting glucose level (mg/dl))/405.

### GTT and ITT

In GTT, mice were intraperitoneally injected with 2 mg/g body weight glucose after overnight (14 h) fasting. In ITT, mice were intraperitoneally injected with insulin (2.0 U/kg) after fasting for 6 hr. Blood samples were collected from the tail tip at various time points (0-120 min), and blood glucose levels were measured using a glucometer (Avenir Biotech, USA). The AUC of GTT or ITT was calculated using the trapezoidal method.

### Western blotting

Proteins were extracted from tissue fragments or cultured cells with lysis buffer containing 10 mM Hepes, pH 7.5, 140 mM NaCl, 10 mM NaF, 1 mM Na_3_VO_4_, 1% Nonidet P-40, 1% sodium deoxycholate, 0.1% SDS, 1 mM PMSF, 1 μg/ml leupeptin, and 1 μg/ml pepstatin A. Protein concentrations were assessed with the BCA protein assay kit (Takara BCA Protein Assay Kit, Takara, Shiga, Japan). SDS-PAGE and Western blotting were performed as previously described (Shintani et al., 20179. Briefly, proteins were separated with SDS-PAGE gels and transferred to PVDF membranes (Immobilon-P, Merck Millipore, USA). Blotted membranes were reacted with the specific primary antibodies shown in Supplementary Table 1 followed by peroxidase-linked secondary antibodies, and proteins were visualized by chemiluminescence using an ECL Reagent (PerkinElmer, USA). Images were acquired with the luminoimage analyzer Luminograph II (ATTO, Tokyo, Japan), and expression levels were normalized and quantified using CS Analyzer4 (ATTO, Tokyo, Japan).

### Collection of Tyr-phosphorylated proteins and separation by 2D electrophoresis

3T3-L1 adipocytes were cultured in medium containing 10 μM AKB9778 and 0.1% DMASO or 0.1% DMSO only for 48 h, and were then retreated with the same fresh medium for 10 min. Proteins were extracted with lysis buffer, and Tyr-phosphorylated proteins were collected using Anti-phosphotyrosine Affinity Beads (Cytoskeleton, USA). Proteins were concentrated using the TCA/acetone precipitation method (Fic et al., 2010), and were then separated by 2D electrophoresis. Isoelectric focusing (IEF) was performed using 7-cm Immobiline DryStrips (pH 3–11 nonlinear) (GE Healthcare Life Sciences, USA) in an Ethan IPGphor II Isoelectric Focusing System (GE Healthcare Life Sciences, USA) according to the manufacturer’s protocol. After IEF, IPG strips were equilibrated in a buffer solution composed of 6 M urea, 2% SDS, 0.375 M Tris-HCl (pH 8.8), 20% glycerol, and 2% DTT for 15 min, followed by a solution of 6 M urea, 2% SDS, 0.375 M Tris-HCl (pH 8.8), 20% glycerol, and 2.5% iodoacetamide for 15 min. 2D SDS-PAGE was performed using 5-10% polyacrylamide gels. Separated proteins were stained with the Silver Stain MS Kit (Fujifilm Wako Pure Chemical Corporation, Osaka, Japan), and silver-stained spots with significant differential expression between AKB9778-treated and control cells were punched out. The liquid-chromatography tandem mass spectrometry/mass spectrometry analysis of samples was entrusted to Nippon Proteomics (Miyagi, Japan).

### Phosphatase assay with synthesized phosphopeptides

Phosphopeptides were synthesized and purified by GenScript Japan (Tokyo, Japan). Phosphatase assays were performed as described previously (Shintani et al., 2006). The amount of phosphate released from phosphopeptides was measured as the malachite green-ammonium molybdate phosphate complex at an absorbance of 595 nm.

### Establishment of stable cell lines expressing WT or mutants of vimentin

The inducible cell lines expressing WT or mutants of vimentin were established by the Tet-One Inducible Expression System (Clontech, USA) according to the manufacturer’s protocol. 3T3-L1 cells were transfected with the pTet-one vector carrying the coding region of WT or mutants of mouse vimentin, which was tagged with the HA epitope at the C terminus, using Lipofectamine LTX (Thermo Fisher Scientific, USA). To isolate cells harboring transfected constructs, cells were grown in medium with 1 μg/ml puromycin. Selected cells were checked for the expression of vimentin proteins with an anti-HA antibody and used for the study. To induce the expression of exogenous WT or mutants of vimentin, cells were exposed to 1 mg/ml doxycycline for the indicated period.

### Protein extraction with NP-40 lysis buffer or SDS lysis buffer

Cellular proteins were sequentially extracted from harvested cells or tissues with equal volumes of NP-40 lysis buffer (10 mM Hepes, pH 7.5, 140 mM NaCl, 10 mM NaF, 1 mM Na_3_VO_4_, 1% Nonidet P-40, 1 mM PMSF, 1 μg/ml leupeptin, and 1 μg/ml pepstatin A), followed by SDS lysis buffer (10 mM Hepes, pH 7.5, 140 mM NaCl, 10 mM NaF, 1 mM Na_3_VO_4_, 0.25% SDS, 1% sodium deoxycholate, 1% Nonidet P-40, 1 mM PMSF, 1 μg/ml leupeptin, and 1 μg/ml pepstatin A).

### Statistical analysis

All assays were performed with 3–8 biological replicates except for phosphatase assays with recombinant proteins, in which three technical replicates were used. The experiment was repeated at least two times except for GTT and ITT, and one representative experiment is shown. Experimenters were blind to information about the samples before obtaining the data. The sample size was calculated from a power analysis using preliminary data obtained in our laboratory under the following assumptions: α = 0.05 and power = 0.8. Data analysis revealed no outliers or other exclusions. Results are expressed as the mean ± standard error of the mean (SEM). Differences between groups were analyzed by the unpaired Student’s *t*-test, a one-way analysis of variance (ANOVA) followed by Tukey’s multiple comparison test, or a two-way ANOVA followed by Tukey’s multiple comparison test using Bell Curve Excel Tokei (SSRI, Japan). Differences were considered to be significant when *P* < 0.05.

## ACKNOWLEDGMENTS

We thank Professor T. Matozaki (Kobe University, Japan) for the kind gift of the anti-PTPRO antibody. We thank M. Goto and K. Wada for their technical assistance, and T. Tanaka and A. Kodama for their secretarial assistance. This work was supported by MEXT/JSPS KAKENHI (Grant number 18K19568 to T. Shintani) and the Japan Health Foundation (to T. Shintani), and in part by MEXT/JSPS KAKENHI (Grant number 19H05659 to M.N.) and Core Research for Evolutional Science and Technology of the Japan Science and Technology Agency (CREST, JST to M.N., grant number JPMJCR1754).

## Data availability

All data generated or analysed during this study are included in the manuscript and Source Data files.

---

**Figure 1-figure supplement 1.**
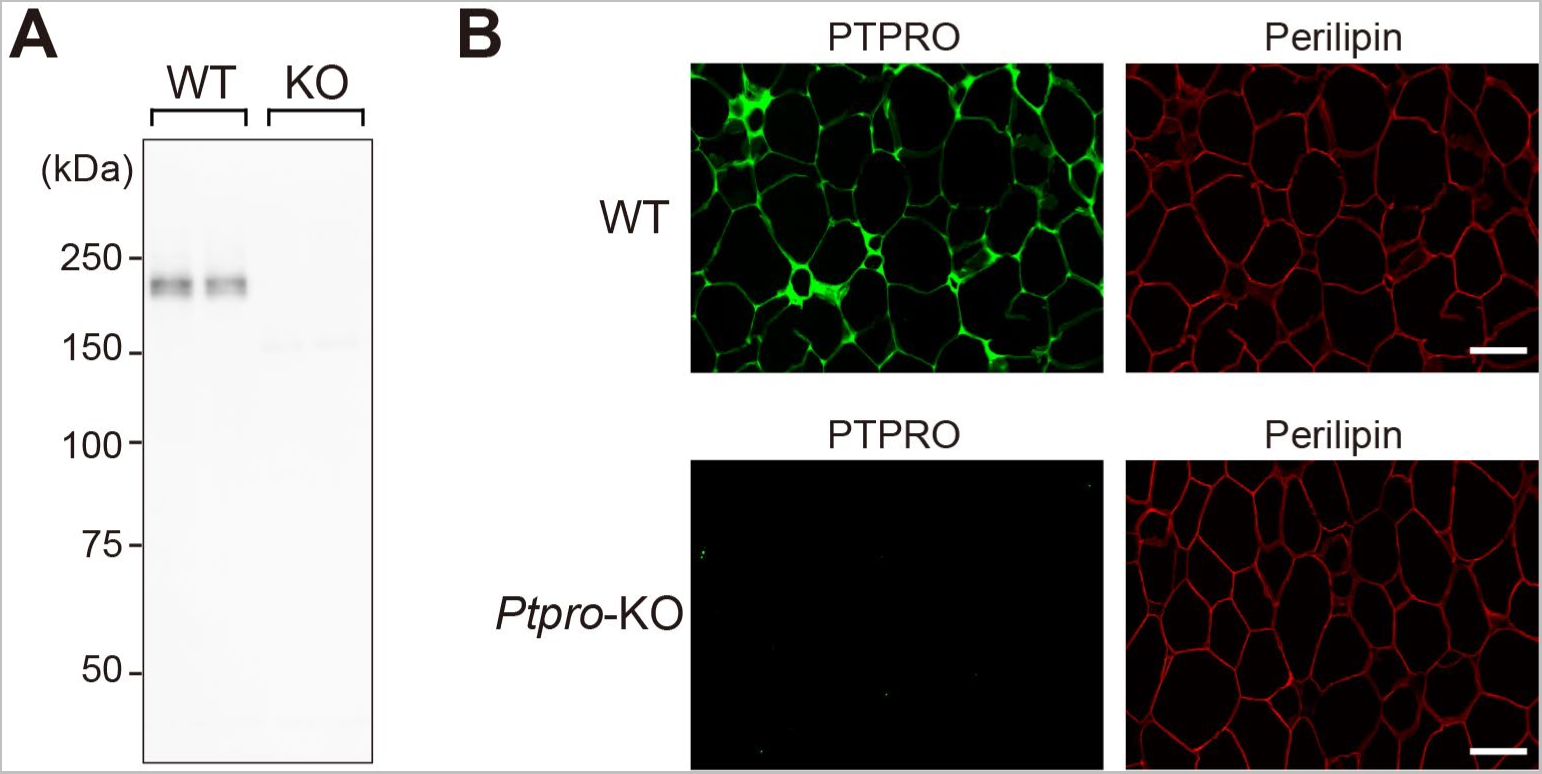
Absence of PTPRO proteins in *Ptpro*-KO mice. (A) Western blot analyses of PTPRO proteins in the epididymal fat lysates of HFHSD-fed WT and *Ptpro*-KO mice at 24 weeks of age. (B) PTPRO immunostaining of the epididymal fat sections of WT and *Ptpro*-KO mice fed HFHSD. Th sections were co-immunostained with antibodies against an adipocyte marker, perilipin. Bars: 100 μm.

**Figure 1-figure supplement 2.**
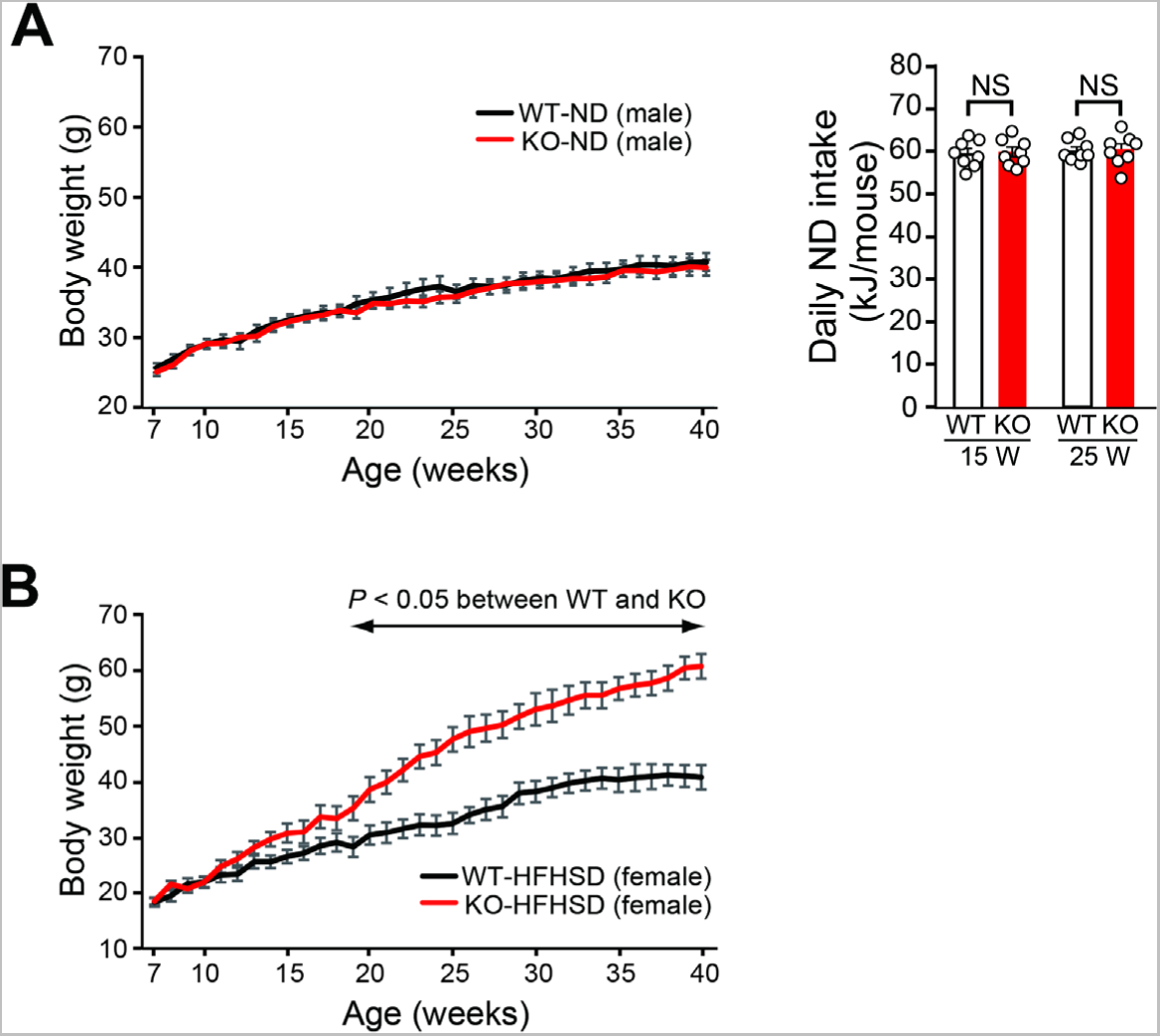
Body weight gain and food intake in *Ptpro*-KO mice fed ND or HFHSD. (A) Weekly body weight gains in wild-type (WT) and *Ptpro*-KO (KO) male mice fed ND (left). Daily ND intake by WT and *Ptpro*-KO male mice at 15 and 25 weeks of age (right). Average food intake is calculated as kJ/mouse. n = 8 each. Values are means ± SEM. *P* values are based on the unpaired Student’s *t*-test. NS, not significant. (B) Weekly body weight gains in WT and *Ptpro*-KO female mice fed HFHSD (n = 8 each). Values are means ± SEM. *P* values are based on the unpaired Student’s *t*-test. **Source data 1.** Body weight gain and food intake in *Ptpro*-KO mice fed ND (Figure 1-figure supplement 2A). **Source data 2.** Body weight gain in *Ptpro-KO* female mice fed HFHSD (Figure 1-figure supplement 2B).

**Figure 1-figure supplement 3.**
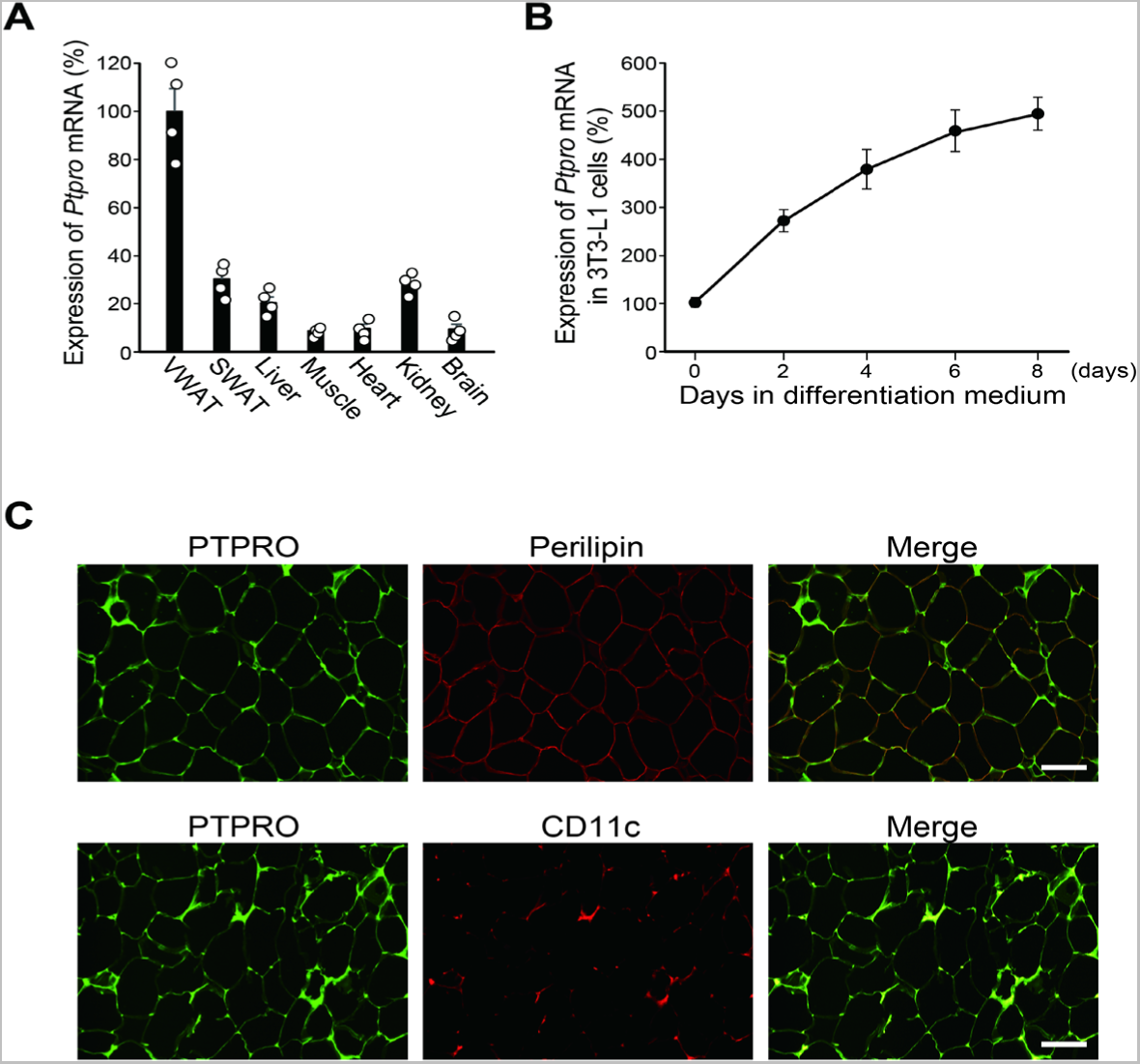
Expression of PTPRO in white adipose tissue. (A) Expression levels of *Ptpro* mRNA in different mouse tissues. Data are normalized to the expression level of *Gapdh* in each tissue and shown as relative values. VWAT, visceral white adipose tissue; SWAT, subcutaneous white adipose tissue. (B) Changes in *Ptpro* mRNA expression in 3T3-L1 adipocytes during maturation. Data are normalized to the expression level of *Gapdh*, and shown as relative values. (C) Immunohistochemical staining of PTPRO in mouse epididymal adipose tissue. Sections are co-stained with antibodies against the adipocyte marker, perilipin (the upper panels), or the M1 macrophage marker, CD11c (the lower panels). Bars: 100 μm. **Source data 1.** Expression levels of *Ptpro* mRNA in different mouse tissues (Figure 1-figure supplement 3A). **Source data 2.** Changes in *Ptpro* mRNA expression in 3T3-L1 adipocytes (Figure 1-figure supplement 3A).

**Figure 1-figure supplement 4.**
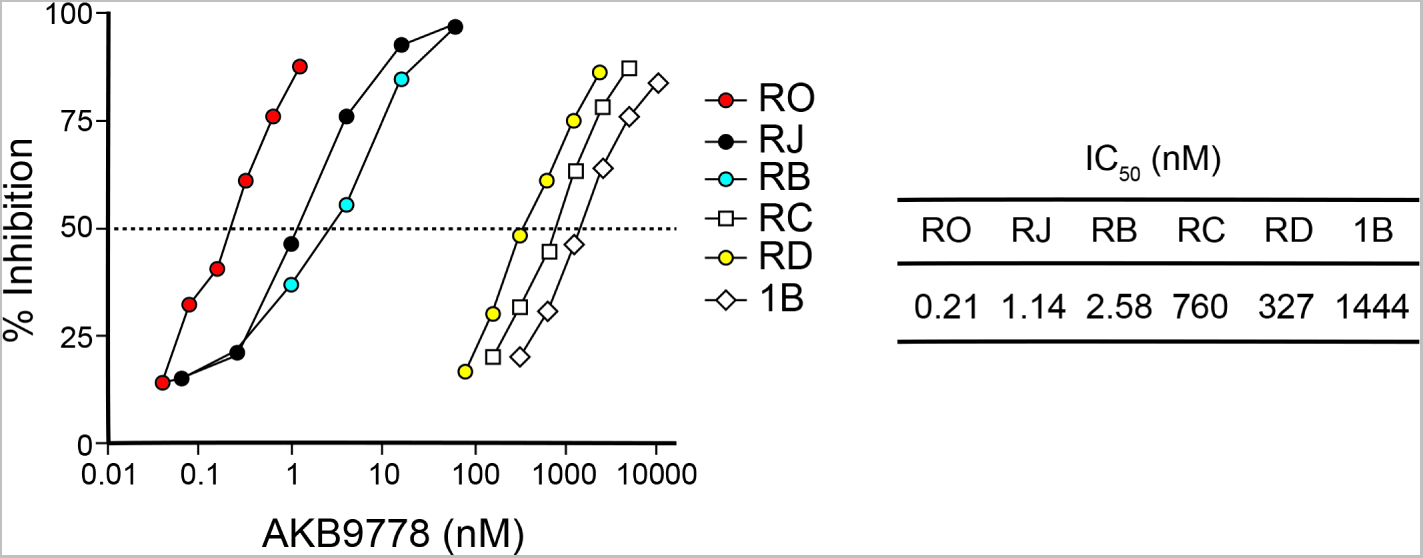
Dose-response curves of the inhibitory activity of AKB9778 against different PTPs. DiFMUP was used as a substrate for assays of the activities of PTPs. Percent inhibition is relative to full activity and no activity controls. Each IC_50_ was calculated based on dose-response curves. **Source data 1.** Inhibitory activity of AKB9778 against different PTPs (Figure 1-figure supplement 4).

**Figure 4-figure supplement 1.**
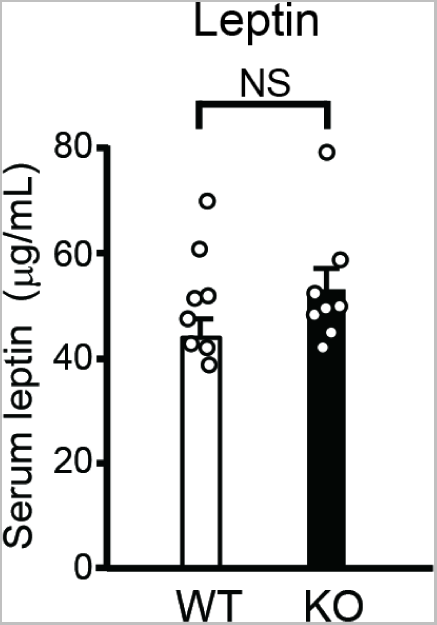
Serum leptin levels in HFHSD-fed WT and *Ptpro*-KO mice. Serum levels of leptin in HFHSD-fed WT and *Ptpro*-KO (KO) mice at 24 weeks of age. n = 8 each. Values are means ± SEM. *P* values are based on the unpaired Student’s *t*-test. NS, not significant (*P* > 0.05). **Source data 1.** Serum leptin levels in HFHSD-fed WT and *Ptpro*-KO mice (Figure 4-figure supplement 1).

**Figure 5-figure supplement 1.**
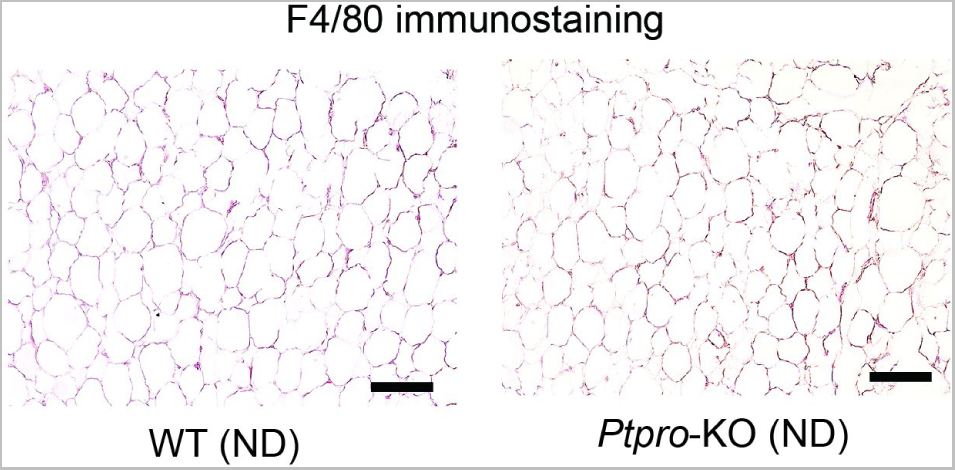
F4/80 immunostaining of the adipose tissue sections of ND-fed WT and *Ptpro*-KO mice. F4/80 immunostaining of the epididymal fat sections of WT and *Ptpro*-KO mice fed ND. Bars: 100 μm.

**Figure 9-figure supplement 1.**
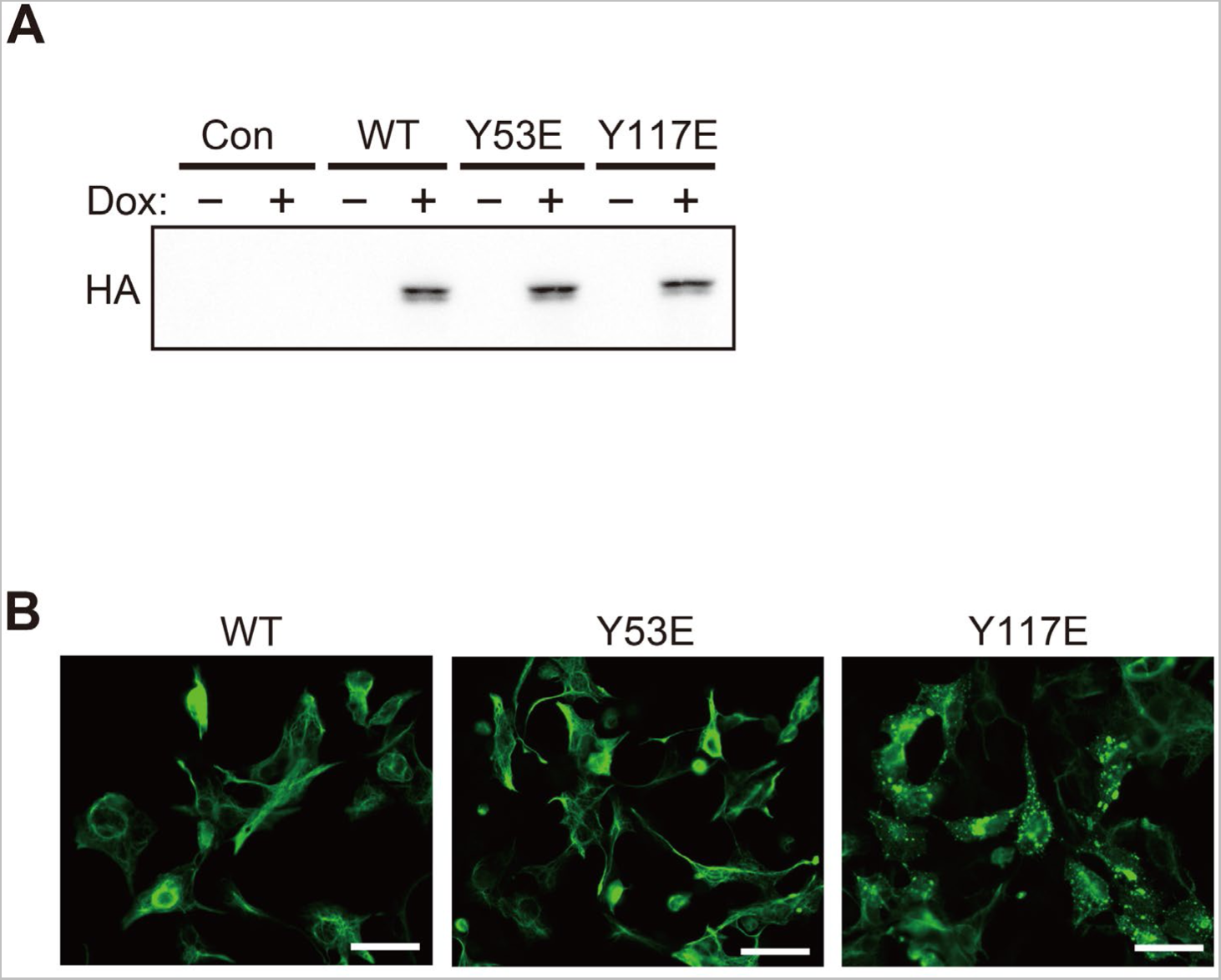
Forced expression of mutant proteins of vimentin in 3T3-L1 adipocytes. (A) Western blot analyses of the WT, Y53E mutant, or Y117E mutant of vimentin expressed in 3T3-L1 adipocytes. All vimentin constructs were tagged with the HA epitope. Cells were treated with or without doxycycline (Dox) for 24 hours to induce expression. Exogeneous and total vimentin proteins were detected with an anti-HA antibody (upper) and anti-vimentin antibody (lower), respectively. (B) Subcellular distributions of exogenous WT and mutants of vimentin expressed in 3T3-L1 adipocytes. Cells were immunostained with an anti-HA antibody. Bars: 50 μm. **Source data 1.** Western blot analyses of the WT, Y53E mutant, or Y117E mutant of vimentin expressed in 3T3-L1 adipocytes (Figure 9-figure supplement 1A).

**Figure 9-figure supplement 2.**
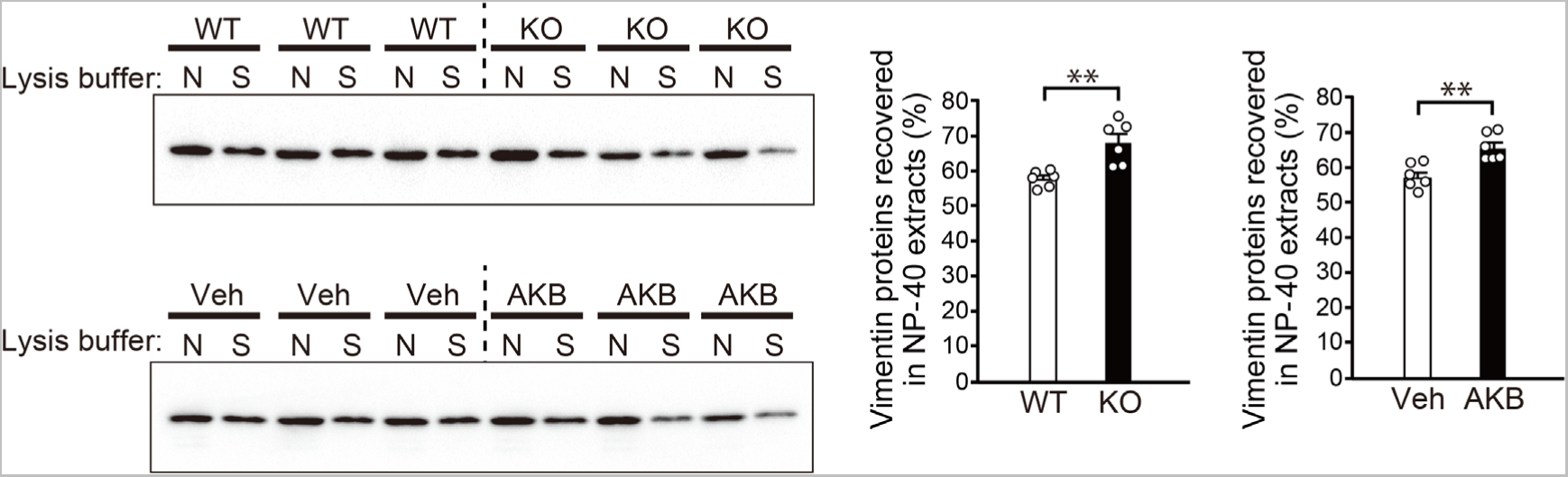
Western blot analyses of vimentin recovered in extracts with NP-40 lysis buffer (N) or SDS lysis buffer (S). Samples were prepared from WT and *Ptpro*-KO (KO) mice fed HFHSD at 24 weeks of age (upper left) and *ob*/*ob* mice treated with vehicle or AKB9778 for 4 weeks from 16 weeks of age (lower left). The middle and right graphs show the results of quantitative analyses (n = 6 each). Values are means ± SEM. *P* values are based on the unpaired Student’s *t*-test. \*\**P* < 0.01. **Source data 1.** Western blot analyses of vimentin recovered in extracts with NP-40 lysis buffer or SDS lysis buffer (Figure 9-figure supplement 2).

